# The lingering effects of Neanderthal introgression on human complex traits

**DOI:** 10.1101/2022.06.07.495223

**Authors:** Xinzhu Wei, Christopher R. Robles, Ali Pazokitoroudi, Andrea Ganna, Alexander Gusev, Arun Durvasula, Steven Gazal, Po-Ru Loh, David Reich, Sriram Sankararaman

**Author notes:** These authors contributed equally (Xinzhu Wei, Christopher R. Robles).

## Abstract

The mutations introduced into the ancestors of modern humans from interbreeding with Neanderthals have been suggested to contribute an unexpected extent to complex human traits. However, testing this hypothesis has been challenging due to the idiosyncratic population genetic properties of introgressed mutations. We developed rigorous methods to assess the contribution of introgressed Neanderthal mutations to heritable trait variation relative to that of modern human variants. We applied these methods to analyze 235,592 introgressed Neanderthal mutations and 96 distinct phenotypes measured in about 300,000 unrelated white British individuals in the UK Biobank. Introgressed Neanderthal mutations have a significant contribution to trait variation consistent with the polygenic architecture of complex phenotypes (contributing 0.1% of heritable variation averaged across phenotypes; *p* = 9.59×10^-9^). However, the contribution of introgressed mutations tends to be significantly depleted relative to modern human mutations matched for allele frequency and linkage disequilibrium (about 57% depletion on average), consistent with purifying selection on introgressed mutations. Different from previous studies (McArthur 2021), we find no evidence for elevated heritability across the phenotypes examined. We identified 348 independent significant associations of introgressed Neanderthal mutations with 64 phenotypes (*p* < 1 ×10^-10^). Previous work (Skov 2021) has suggested that a majority of such associations are likely driven by statistical association with nearby modern human variants that are the true causal variants. We therefore developed a customized statistical fine-mapping methodology for introgressed mutations that led us to identify 112 regions (at a false discovery proportion of 16%) across 47 phenotypes containing 4,303 unique genetic variants where introgressed mutations are highly likely to have a phenotypic effect. Examination of these mutations reveal their substantial impact on genes that are important for the immune system, development, and metabolism. Our results provide the first rigorous basis for understanding how Neanderthal introgression modulates complex trait variation in present-day humans.

## Introduction

Genomic analyses have revealed that present-day non-African human populations inherit 1-4% of their genetic ancestry from introgression with Neanderthals (Green 2010, Prüfer 2014). This introgression event introduced uniquely Neanderthal variants into the ancestral out-of-Africa human gene pool, which may have helped this bottleneck population survive the new environments they encountered (Mendez 2012, Abi-Rached 2011, Sankararaman 2014, Vernot 2014, Racimo 2015, Gittelman 2016). On the other hand, the bulk of Neanderthal variants appear to have been deleterious in the modern human genetic background leading to a reduction in Neanderthal ancestry in conserved genomic regions (Sankararaman 2014, Vernot 2014, Harris 2016, Juric 2016, Petr 2019). Systematically studying these variants can provide insights into the biological differences between Neanderthals and modern humans and the evolution of human phenotypes in the 50,000 years since introgression.

In principle, studying Neanderthal-derived mutations in large cohorts of individuals measured for diverse phenotypes can help understand the biological impact of Neanderthal introgression. Previously, Dannemann and Kelso (Dannemann 2017) showed that some Neanderthal introgressed variants are significantly associated with traits such as skin tone, hair color, and height based on Genome-Wide Association Studies (GWAS) in British samples. However, using data from Iceland, Skov et al. (Skov 2020) found that most of the significantly associated Neanderthal introgressed SNPs are in the proximity of strongly associated non-archaic variants. They suggested that these associations at Neanderthal introgressed SNPs were driven by the associations at linked non-archaic variants, indicating a limited contribution to modern human phenotypes from Neanderthal introgression. In contrast to these attempts to associate individual introgressed variants with a trait, studies have attempted to measure the aggregate contribution of introgressed Neanderthal SNPs to trait variation (Simonti 2016, McArthur 2021). A recent study by McArthur and colleagues (McArthur 2021) estimated the proportion of heritable variation that can be attributed to introgressed variants though their approach is restricted to common variants (minor allele frequency > 5%) that represent a minority of introgressed variants. Despite these attempts, assessing the contribution of introgressed Neanderthal variants towards specific phenotypes remains challenging. The first challenge is that variants introgressed from Neanderthals are rare on average (due to the low proportion of Neanderthal ancestry in present-day genomes). The second challenge arises from the unique evolutionary history of introgressed Neanderthal variants resulting in distinct population genetic properties at these variants which can, in turn, confound attempts to characterize their effects. As a result, attempts to characterize the systematic impact of introgressed variants on complex phenotypes need to be rigorously assessed.

To enable analyses of genome-wide introgressed Neanderthal variants in large sample sizes, we selected and added Single Nucleotide Polymorphism (SNPs) that tag introgressed Neanderthal variants to the UKBiobank Axiom Array that was used to genotype the great majority of the approximately 500,000 individuals in the UK Biobank (UKBB) (Bycroft 2018). We used a previously compiled map of Neanderthal haplotypes in the 1000 Genomes European populations (Sankararaman 2014) to identify introgressed SNPs that tag these haplotypes. After removing SNPs that are well-tagged by those previously present on the UK Biobank (UKBB) array, we used a greedy algorithm to select 6,027 SNPs that tag the remaining set of introgressed SNPs at *r*^2^ > 0. 8 which were then added to the UKBB genotyping array to better tag Neanderthal ancestry. These SNPs allow variants of Neanderthal ancestry to be confidently imputed and allow us to identify a list of 235,592 mutations that are likely to be Neanderthal-derived (termed Neanderthal Informative Mutations or NIMs) out of a total of 7,774,235 QC-ed SNPs in UKBB (see **Methods; Note S1**).

The goals of our study are threefold: 1) to estimate the contribution of NIMs to phenotypic variation in modern humans, 2) to test the null hypothesis that a NIM has the same contribution to phenotypic variation as a non-introgressed modern human SNP, and 3) to pinpoint regions of the genome at which NIMs are highly likely to modulate phenotypic variation. We develop rigorous methodology for each of these goals which we validate in simulations. We then applied these methods to 96 distinct phenotypes measured in about 300,000 unrelated white British individuals in UKBB.

## Results

### The contribution of Neanderthal introgressed variants to trait heritability

To understand the contribution of Neanderthal introgressed variants to trait variation, we aim to estimate the proportion of phenotypic variance attributed to NIMs (NIM heritability) and to test the null hypothesis that per-NIM heritability is the same as the heritability of a non-introgressed modern human (MH) SNP. We first annotated each of the 7,774,235 QC-ed SNPs in UKBB as either a NIM or a MH SNP (see **Methods**). NIMs include SNPs created by mutations which likely originated in the Neanderthal lineage after the human-Neanderthal split. SNPs that are not defined as NIMs are annotated as MH SNPs which likely originated in the modern human lineage or the human-Neanderthal common ancestor.

To estimate NIM heritability, we used a recently proposed method (RHE-mc) that can partition the heritability of a phenotype measured in large samples across various genomic annotations (Pazokitoroudi 2020). We applied RHE-mc with genomic annotations that correspond to the ancestry of each SNP (NIM vs MH) to estimate NIM heritability (*h*^2^_NIM_). We also attempted to estimate whether per-NIM heritability is the same as the per-SNP heritability of MH SNPs (Δ_*h*^2^_). A positive (negative) value of Δ 2 indicates that, on average, a NIM makes a larger (smaller) contribution to phenotypic variation relative to a MH SNP.

To assess the accuracy of this approach, we performed simulations where NIMs are neither enriched nor depleted in heritability (true Δ_*h*^2^_ = 0). Following previous studies of the genetic architecture of complex traits (Evans 2018, Gazal 2018), we simulated phenotypes (across 291,273 unrelated white British individuals and 7,774,235 SNPs) with different architectures where we varied heritability, polygenicity, and how the effect size at a SNP is coupled to its population genetic properties (the minor allele frequency or MAF at the SNP and the linkage disequilibrium or LD around a SNP). We explored different forms of MAF-LD coupling where BASELINE assumes that SNPs with phenotypic effects are chosen randomly, RARE (COMMON) assumes that rare (common) variants are enriched for phenotypic effects, and HIGH (LOW) assumes that SNPs with high (low) levels of LD (as measured by the LD score (Finucane 2015)) are enriched for phenotypic effects (see **Methods**). Estimates of 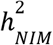 and Δ_*h*^2^_ tend to be miscalibrated (**Fig. 1ab**). The miscalibration is particularly severe when testing Δ _2_ so that a test of the null hypothesis has a false positive rate of 0.55 across all simulations (at a p-value threshold of 0.05).

**Figure 1.**
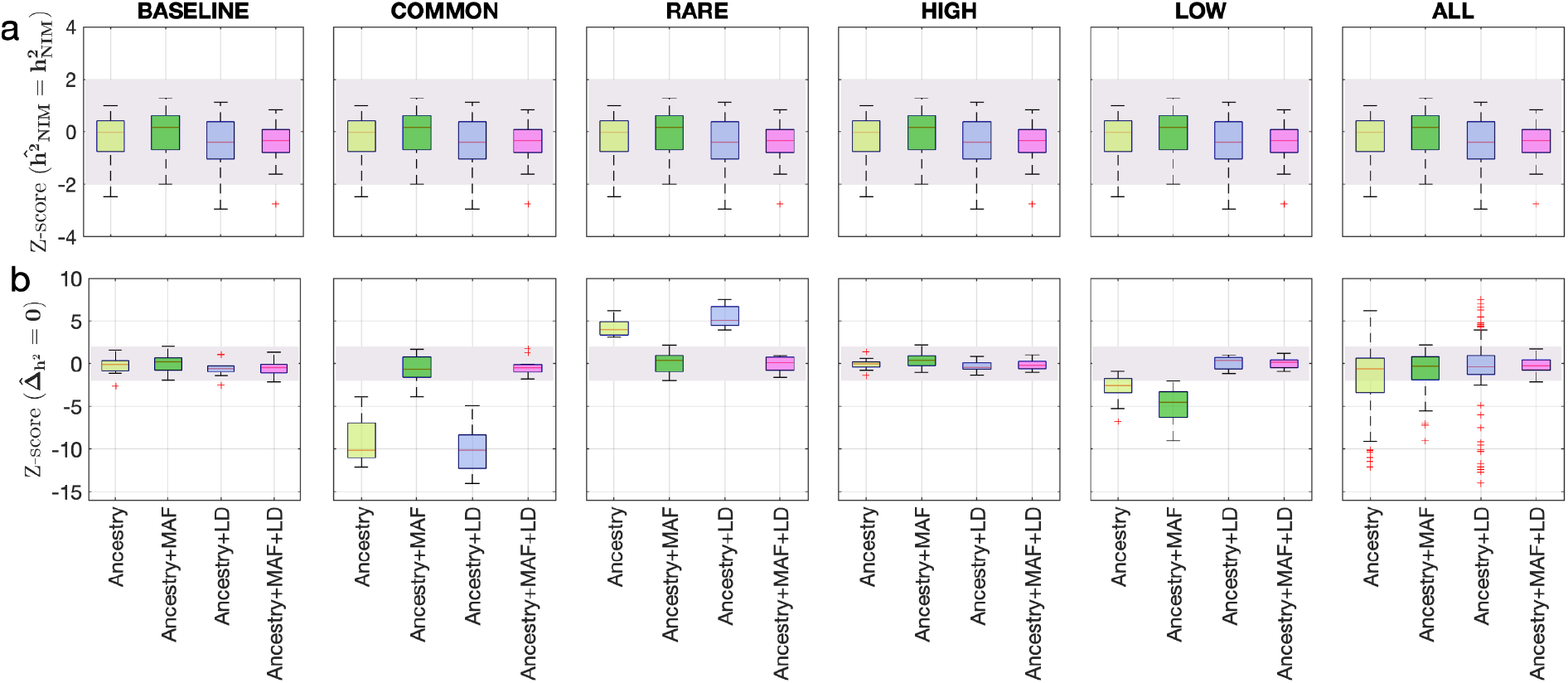
Benchmarking approaches for estimating the heritability components of Neanderthal introgression. We group simulations by relationships between minor allele frequency (MAF) and local linkage disequilibrium at a SNP on effect size (MAF-LD coupling): BASELINE, COMMON, RARE, HIGH, LOW. In each group, we perform 12 simulations with varying polygenicity and heritability (see **Methods**). Additionally, we combine results from all simulations together as ALL. We plot the distributions of two Z-scores (y-axis), one on each row: **(a)** Z-score 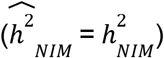 tests whether the estimated and true NIM heritability are equal, and **(b)** Z -score 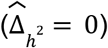 tests whether the estimated per-NIM heritability is the same as the per-SNP heritability of MH SNPs (see **Methods**). In each panel, we present results from a variance components analysis method (RHE-mc) using four different input annotations: ancestry only where ancestry is either NIM or MH, ancestry + MAF, ancestry + LD, ancestry + MAF + LD. A calibrated method is expected to have Z-scores distributed around zero and within ±2 (shaded region). Among all tested approaches, only RHE-mc with ancestry + MAF + LD annotations is calibrated across simulations.

To understand these observations, we compared the Minor Allele Frequencies (MAF) and LD scores at NIMs to MH SNPs. We observe that NIMs tend to have lower MAF **(Fig.2a)** and higher LD scores compared to MH SNPs **(Fig. 2b)** (the average MAF of NIMs and MH SNPs are 3.9% and 9.9%, respectively while their average LD scores are 170.6 and 64.9). Among the QC-ed SNPs, 76.9% of NIMs have MAF > 1%, and 27.7% have MAF > 5%, in contrast to 61.6% and 41.6% of MH SNPs. Distinct from MH SNPs, the MAF and LD score of NIMs tend not to increase with each other **(Fig. 2cd)**.

**Figure 2.**
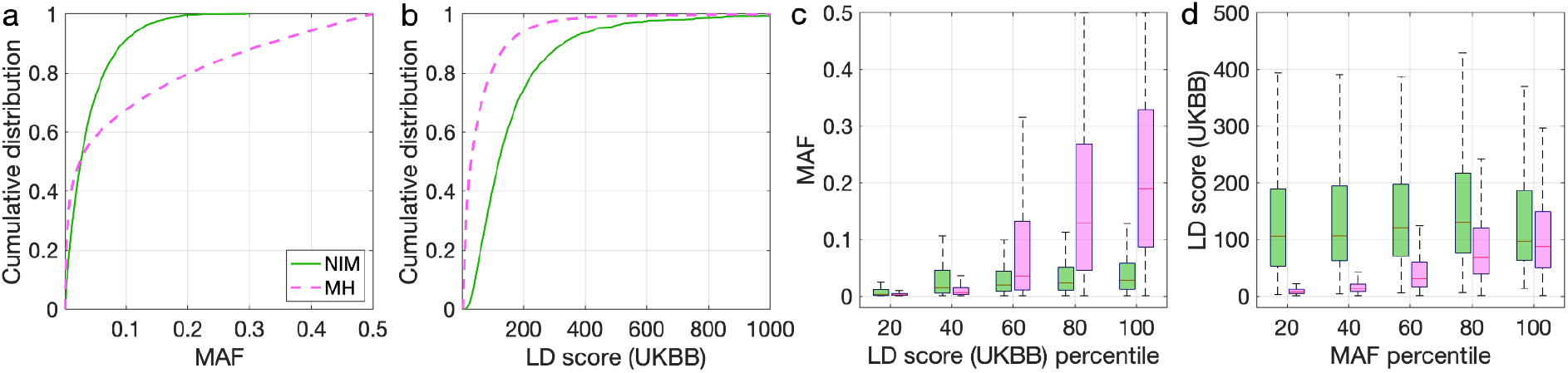
Distributions of minor allele frequency (MAF) and LD-score in NIMs and MH SNPs. Empirical cumulative distribution functions of **(a)** MAF and **(b)** LD scores of NIMs (in solid green line) and MH SNPs (in pink dashed line) estimated in the UK Biobank (UKBB). **(c)** Boxplots of MAFs of NIMs (on the left filled in green) and MH SNPs (on the right side filled in pink) while controlling for LD score (UKBB). **(d)** Boxplots of LD score (UKBB) of NIMs and MH SNPs while controlling for MAF. NIMs and MH SNPs are divided by the 20, 40, 60, 80, 100 **(c)** LD score (UKBB) percentile or MAF percentile **(d)** based on all QC-ed SNPs (7,774,235 imputed SNPs with MAF > 0.001). The lower and upper edges of a box represent the first and third quartile (qu1 and qu3), respectively; the horizontal red line inside the box indicates median (md); the whiskers extend to the most extreme values inside inner fences, md ± 1.5 (qu3–qu1).

To account for the differences in the MAF and LD scores across NIMs and MH SNPs, we applied RHE-mc with annotations corresponding to the MAF and the LD score at each SNP (in addition to the ancestry annotation that classifies SNPs as NIM vs. MH) to estimate NIM heritability (*h*^2^_*NIM*_) and to test whether per-NIM heritability is the same as the per-SNP heritability of MH SNPs i.e., Δ_*h*_2__ = 0 (see **Methods, Note S4**). Our simulations show that RHE-mc with SNPs assigned to annotations that account for both MAF and LD (in addition to the ancestry annotation that classifies SNPs as NIM vs. MH) is accurate both in the estimates of 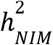 (**Fig. 1a**) and in testing the null hypothesis that Δ _2_ = 0 (the false positive rate of a test of Δ_*h*_2__ = 0 is 0.017 at a p-value threshold of 0.05; **Fig. 1b**). On the other hand, not accounting for either MAF or LD leads to poor calibration (**Fig. 1;** we observe qualitatively similar results when estimating genome-wide SNP heritability; **Fig S1**).

We then applied RHE-mc with ancestry+MAF+LD annotations to analyze a total of 96 UKBB phenotypes that span 14 broad categories (**Data S2**). In all our analyses, we include the top five PCs estimated from NIMs (NIM PCs) as covariates in addition to the top twenty genetic PCs estimated from common SNPs, sex, and age (see **Methods**). The inclusion of NIM PCs is intended to account for stratification at NIMs that may not be adequately corrected by including genotypic PCs estimated from common SNPs (we also report concordant results from our analyses when excluding NIM PCs; **Note S3** and **Fig. S3–S4**).

We first examined NIM heritability to find six phenotypes with significant NIM heritability (Z-score 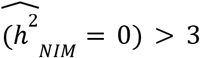): body fat percentage, trunk fat percentage, whole body fat mass, overall health rating, gamma glutamyltransferase (a measure of liver function), and forced vital capacity (FVC) (**Fig. 3ac**). Meta-analyzing within nine categories that contain at least four phenotypes, we find that *meta* – 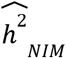 is significantly larger than zero for anthropometry, blood biochemistry, bone densitometry, kidney, liver, and lung but not for blood pressure, eye, lipid metabolism (*p* < 0.05 accounting for the number of hypotheses tested). Meta-analyzing across all phenotypes with low correlation, we obtain overall NIM heritability estimates 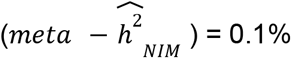 (one-sided *p*= 9.59×10^-9^). The estimates of NIM heritability are modest as would be expected from traits that are highly polygenic and given that NIMs account for a small percentage of all SNPs in the genome (see **Methods**).

**Figure 3.**
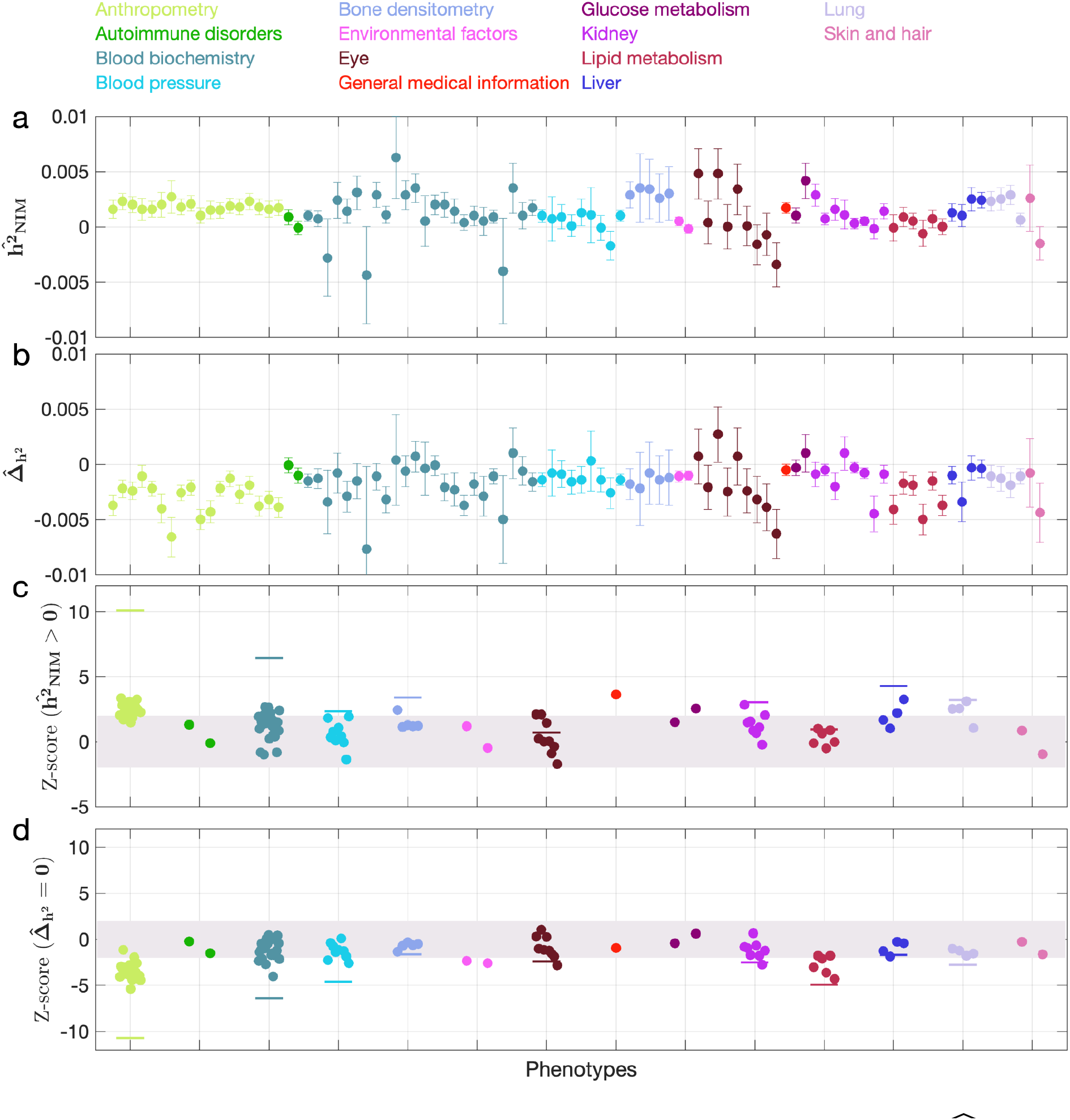
NIM heritability in UKBB phenotypes. **(a)** Estimates of NIM heritability 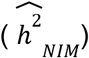 and **(c)** the Z-score of 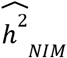 (testing the hypothesis that NIM heritability is positive) for each UKBB phenotype. Analogously, **(b)** estimates of 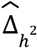 and Z-score **(d)** of 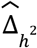 (testing the hypothesis that per-NIM heritability is equal to per-SNP heritability at MH SNPs after controlling for MAF and LD). Phenotypic categories are shown in alphabetical order and listed on the top of panel **(a)** in the same color and alphabetical order (from top to bottom, and left to right) as they are in the figure. The estimate for each phenotype is shown as one colored dot, on the x-axis based on its phenotypic category, and on the y-axes based on its Z-score 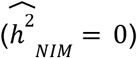 and Z-score 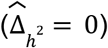, for panels **(c)** and **(d)** respectively. For each phenotypic category with at least four phenotypes, their Z-scores from random effect meta-analysis are plotted with the flat colored lines (see Methods). The color shades cover Z-scores around zero and within ±2.

We next tested whether the average heritability at a NIM is larger or smaller compared to a MH SNP 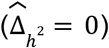. We find seventeen phenotypes with significant evidence of depleted NIM heritability that include standing height, body mass index, and HDL cholesterol (Z-score < −3; **Fig. 3bd**). Five phenotypic categories show significant NIM heritability depletion (anthropometry, blood biochemistry, blood pressure, lipid metabolism, lung) in meta-analysis. Meta-analyzing across phenotypes, we find a significant depletion in NIM heritability (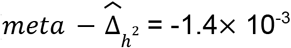, *p*= 2.55×10^-11^). On average, we find that heritability at NIMs is reduced by about 57% relative to a modern human variant with matched MAF and LD characteristics. In contrast to the evidence for depletion in NIM heritability, we find no evidence for traits with elevated NIM heritability across the phenotypes analyzed. Despite the observation that NIMs have been primarily under purifying selection for thousands of generations (Harris and Nielsen 2016, Petr 2019), they still make a substantial contribution to phenotypic variation in present-day humans.

Finally, we investigated the impact of controlling for MAF and LD on our findings in UKBB. Analyses that do not control for MAF and LD tend to broadly correlate with our results that control for both (Pearson’s *r* = 0.96, 0.68, and 0.65 and *p* < 10^-12^ among 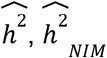, and 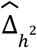). However, these analyses underestimate both heritability (**Fig. 4a**) and NIM heritability (**Fig. 4b**), resulting in apparent NIM heritability depletion (Z-score < −3) in 83 of the 96 phenotypes (**Fig. 4c**). While yielding qualitatively similar conclusions about the depletion in heritability at NIMs relative to MH SNPs, prior knowledge that per SNP heritability of complex traits can be MAF and LD dependent (Evans 2018) coupled with our extensive simulations lead us to conclude that controlling for MAF and LD lead to more accurate results.

**Figure 4.**
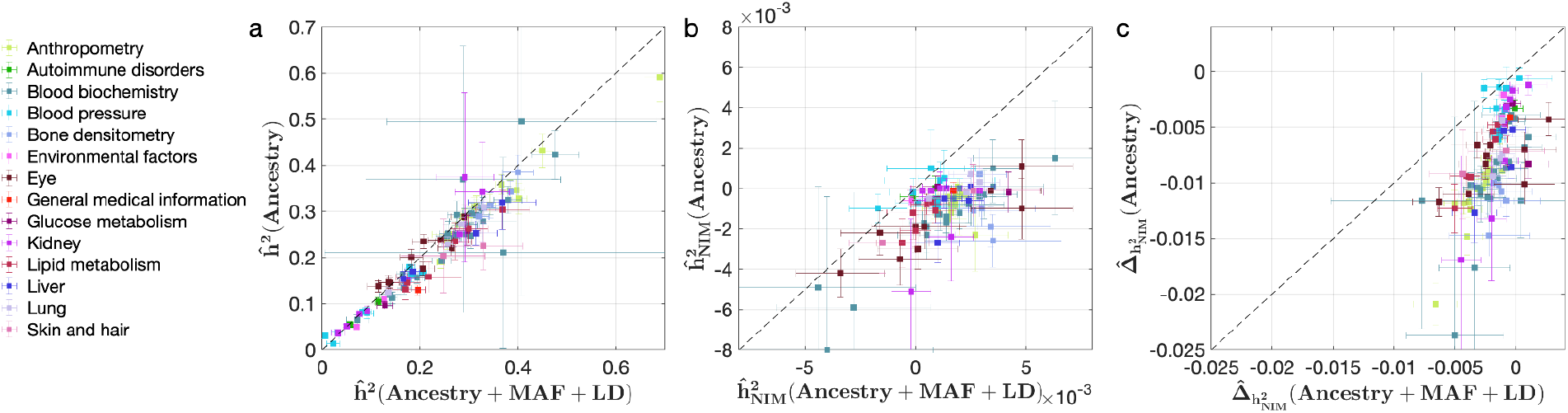
Comparing heritability analyses with and without controlling for MAF and LD in UKBB phenotypes. Each phenotype is shown with one dot colored by the phenotypic category it belongs to, on the y-axis based on its point estimate and standard error (estimated by RHE-mc with Ancestry annotation) and on the x-axis based on its point estimate and standard error (estimated by RHE-mc with ancestry + MAF + LD annotation). Estimates shown are **(a)** total heritability 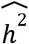, **(b)** NIM heritability 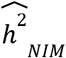, and **(c)** the difference between per-NIM heritability and matched MH SNPs heritability 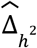. Not controlling for MAF and LD leads to underestimation of NIM heritability, which leads to false positives when testing whether heritability at a NIM is elevated or depleted relative to a MH SNP.

### Identifying genomic regions at which introgressed variants influence phenotypes

Having documented an overall contribution of NIMs to phenotypic variation, we focus on identifying individual introgressed variants that modulate variation in complex traits. We first tested individual NIMs for association with each of 96 phenotypes (controlling for age, sex, twenty genetic PCs (estimated from common SNPs), and five NIM PCs (that account for potential stratification that is unique to NIMs). We obtained a total of 13,075 significant NIM-phenotype associations in 64 phenotypes with 8,018 unique NIMs (*p* < 10^-10^ that accounts for the number of SNPs and phenotypes tested) from which we obtain 348 significant NIM-phenotype associations with 294 unique NIMs after clumping associated NIMs by LD (see **Methods**).

A limitation of the association testing approach is the possibility that a NIM might appear to be associated with a phenotype simply due to being in LD with a non-introgressed variant (Skov 2020). We formally assessed this approach in simulations of phenotypes with diverse genetic architectures described previously where the identities of causal SNPs are known. A NIM that was found to be associated with a phenotype (*p* < 10^-10^) was declared a true positive if the 200 kb region surrounding the associated NIM contains any NIM with a non-zero effect on the phenotype and a false positive otherwise. Averaging across all genetic architectures, the False Discovery Proportion (FDP; the fraction of false positives among the significant NIMs) of the association testing approach is around 30% **(Fig. 5b)**. Hence, finding NIMs that are significantly associated with a phenotype does not confidently localize regions at which introgressed variants affect phenotypes.

**Figure 5.**
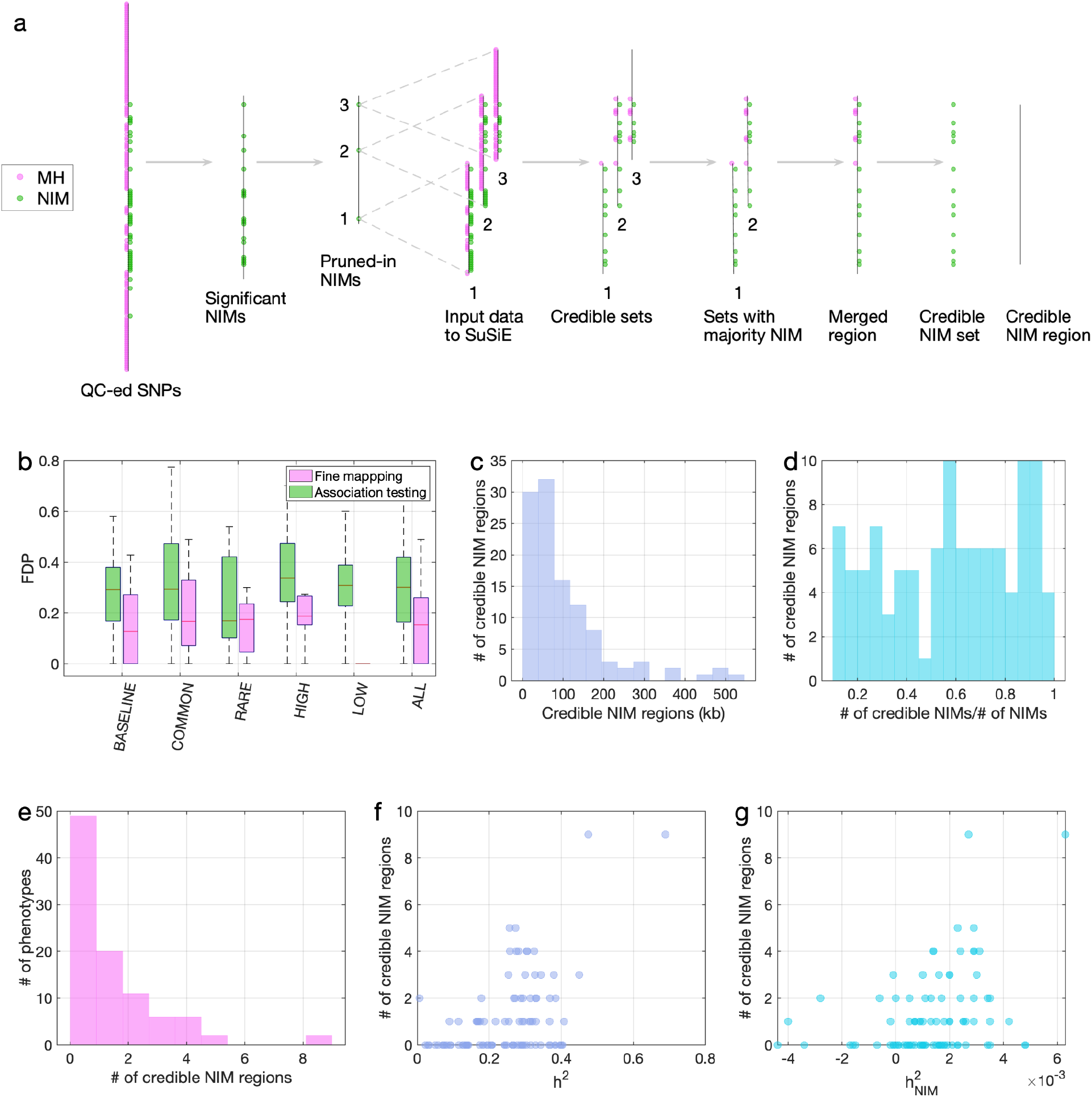
Fine mapping of NIMs in simulations and the UKBB. **(a)** Fine mapping pipeline to identify NIMs that aims to identify genomic regions at which NIMs are likely to modulate phenotypic variation (credible NIM regions). **(b)** Comparison of approaches for identifying credible NIM regions. For each simulation, False Discovery Proportion (FDP) is computed for association testing compared to our pipeline (combining association testing and fine-mapping). The distributions of the FDP are shown across genetic architectures (summarized across groupings of coupling of effect size, MAF and LD) and summarized across architectures (ALL). Our approach to identifying credible NIMs decreases FDP in all studied architectures (the LOW LD setting has a median and quartiles of zero across replicates). **(c)** The distribution of the length of credible NIM regions across 96 UKBB phenotypes. **(d)** Distribution of the ratio between the number of credible NIMs and number of tested NIMs (in the example of panel **(a)**, the number of tested NIMs is the union of NIMs in input to the fine-mapping software (SuSiE) 1 and 2). This figure shows that our fine mapping approach is effective in prioritizing NIMs that affect phenotype. **(e)** The distribution of the number of credible NIM regions among phenotypes. The number of credible NIM regions is positively correlated with **(f)** heritability **(g)** NIM heritability.

To improve our ability to identify NIMs that truly modulate phenotype, we designed a customized pipeline that combines association testing with a fine-mapping approach that integrates over the uncertainty in the identities of causal SNPs to identify sets of NIMs that plausibly explain the association signals at a region (**Fig. 5a)**. Our pipeline starts with a subset of significantly associated NIMs that are relatively independent *(p* < 10^-10^) followed by the application of a statistical fine-mapping method (SuSiE) within the 200kb window around each NIM signal (Wang 2020) and additional post-processing to obtain a set of NIMs that have an increased probability of being causal for a trait. We term the NIMs within this set **credible NIMs** while the shortest region that contains all credible NIMs in a credible set is termed the **credible NIM region** (see **Methods**; **Fig. 5a**).

We employed the same simulations as previously described to evaluate our fine-mapping approach. The fine mapping approach yields a reduction in the FDP relative to association mapping (FDP of 15.6% on average; **Fig. 5b**) while attributing the causal effect to a few dozen NIMs within the credible NIM set (mean: 79, median: 54 NIMs across all simulations). Applying our pipeline to the set of 96 UKBB phenotypes, we identified a total of 112 credible NIM regions containing 4,303 unique credible NIMs across 47 phenotypes (**Fig. 6a**). The median length of credible NIM regions, 65.7kb (95% CI: [4.41kb, 469.3 kb]) is close to the expected length of Neanderthal introgressed segments (Skov 2020) suggesting that the resolution of our approach is that of an introgressed LD block (**Fig. 5c**). While fine mapping generally attributes the causal signal to a subset of the tested NIMs (mean: 55.8, median: 37 NIMs across phenotypes), the degree of this reduction varies across regions likely reflecting differences in the LD among NIMs (**Fig. 5d**). We do not detect any credible NIM in 49 out of 96 phenotypes potentially due to the limited power of our procedure that aims to control the FDR (**Fig. 5e**). The sensitivity of our method is affected by both total heritability (**Fig. 5f**, Pearson’s *r* = 0.49, *p* = 3.3×10^-7^) and NIM heritability (**Fig. 5g**, Pearson’s *r* = 0.36, *p* = 3.3×10^-4^). A linear model that uses both total heritability and NIM heritability to predict the number of credible sets yields *r*^2^ = 0.29, *p* = 1.3× 10^-5^ and 0.015, respectively), while linear models with only total heritability or only NIM heritability result in statistically lower *r*^2^ (0.24 and 0.13, respectively).

**Figure 6.**
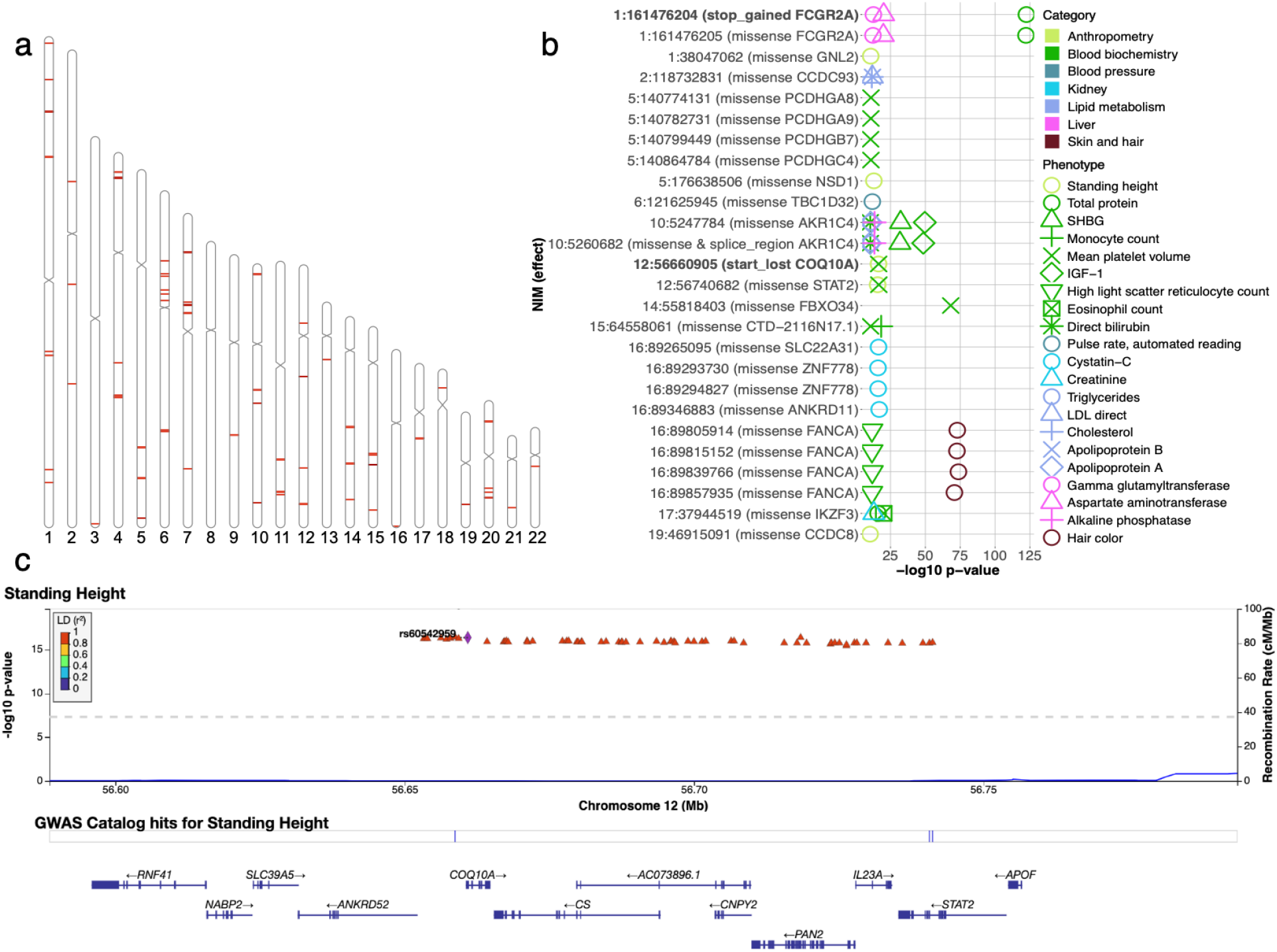
Analysis of credible NIMs. **(a)** Distribution of credible NIMs across the genome **(b)** High and moderate impact credible NIMs annotated by SnpEff software (Cingolani 2012). A total of 26 credible NIMs have high (marked in bold) or moderate impact effects on nearby genes (chromosome number and hg19 coordinates). The effect of the SNP and the gene name are displayed. This plot shows significant associations of these NIMs with specific phenotypes (color denotes the phenotype category). **(c)** Plot of 300kb region surrounding rs60542959 (marked in black diamond; hg19 coordinates), a credible NIM for standing height that results in loss of the start codon in COQ10A. The plot displays other significantly associated NIMs in the region along with their LD (r^2^) to rs60542969 in 1000 Genomes Europeans (Boughton 2021).

### Examination of the functional impact of credible NIMs

We annotated all 4,303 unique credible NIMs using SnpEff (Cingolani 2012) to identify a total of 26 NIMs with high (e.g., start codon loss, stop codon gain) or moderate impact (nonsynonymous variants) on genes (**Fig. 6b, SI Data S7**). We identified two credible NIMs, rs9427397 (1:161,476,204 C>T) and rs60542959 (12:56,660,905 G>T), that have a high impact on protein sequences. The 1:161,476,204 C>T mutation, a NIM that is associated with increased gamma glutamyltransferase and aspartate aminotransferase (enzymes associated with liver function) and decreased total protein levels in blood, introduces a premature stop codon in the FCGR2A gene (**Fig. S7**). FCGR2A codes for a receptor in many immune cells, such as macrophages and neutrophils, and is involved in the process of phagocytosis and clearing of immune complexes. This NIM is in a region that contains SNPs shown in several GWAS linked to rheumatoid arthritis (Okada 2014, Laufer 2019). The other high impact mutation, 12:56,660,905 G>T (rs60542959), results in the loss of the start codon in COQ10A, and this SNP is a credible NIM for both mean platelet volume and standing height (**Fig. 6c**). COQ10 genes (A and B) are important in respiratory chain reactions. Deficiencies of CoQ10 (MIM 607426) have been associated with encephalomyopathy, infantile multisystemic disease; cerebellar ataxia, and pure myopathy (Quinzii 2009). The start codon in COQ10A is conserved among mammals with its loss having a potentially significant effect on COQ10A expression in immune cells (Kubota 2020).

In addition, we detect 24 credible NIMs that function as missense mutations in 19 genes. Seven out of the 19 genes are known to have immune related functions (FCGR2A, PCDHG (A8, A9, B7, C4), STAT2, and IKZF3). The NIM in STAT2 (rs2066807, 12:56,740,682 C>G) was the first adaptive introgression locus to be identified (Mendez 2012). The STAT2 introgressed variant segregates at 0.066 frequency in the UKBB white British and leads to an I594M amino acid change in the corresponding protein. STAT2 gene and COQ10A are neighboring genes thereby providing an example of an introgressed region that potentially impacts function at multiple genes (**Fig. 6c**).

At least seven of the 12 genes not known to be immune related have other important functions documented in the literature, such as DNA replication/damage (FANCA, CCDC8), transition in meiosis (FBXO34), detoxification/metabolism (AKR1C4), and neurological/developmental (ZNF778, ANKRD11, TBC1D32) functions. rs17134592 (10:5260682 C>G) is a non-synonymous mutation in AKR1C4, a gene that is involved in the metabolism of ketone-containing steroids in the liver. The NIM is associated with increased serum bilirubin levels (*p* = 3×10^-11^) **(Fig. S7a)** while also being associated with increased levels of alkaline phosphatase, insulin-like growth factor 1 (IGF1) and decreased apolipoprotein A, sex hormone binding globulin (SHBG) and triglyceride levels. rs17134592 has been identified to be a splicing QTL that is active in the liver and testis in the GTeX data **(Fig. S7b)**. This NIM alters Leucine to Valine (L311V) which, in combination with the tightly-linked non-synonymous variant rs3829125 (S145C) in the same gene, have been shown to confer a three-to-five-fold reduction in catalytic activity of the corresponding enzyme (3-alpha hydroxysteroid dehydrogenase) in human liver (Kume 1999). Interestingly, the single amino acid change S145C did not significantly alter enzyme activity suggesting the importance of the amino acid residue at position 311 for the substrate binding of the enzyme.

## Discussion

Our analysis demonstrates the complex influence of Neanderthal introgression on complex human phenotypes. The assessment of the overall contribution of introgressed Neanderthal alleles to phenotypic variation indicates a pattern where, taken as a group, these alleles tend to be depleted in their impact on phenotypic variation (with about a third of the studied phenotypes showing evidence of depletion). This pattern is consistent with these alleles having entered the modern human population roughly 50,000 years ago and being subject to purifying selection. Selection to purify deleterious introgressed variants, coupled with stabilizing selection on human complex traits, could result in introgressed heritability depletion such that the remaining introgressed variants in present-day humans tend to have smaller phenotypic effects compared to other modern human variants.

Nevertheless, we document a modest but significant contribution of introgressed alleles to variation in a number of phenotypes. In contrast to the previous heritability analyses by McArthur et al. (McArthur 2020), we did not find any NIM heritability enrichment in the 96 phenotypes. This discrepancy could be due to the different methods and NIMs used in the two studies. McArthur et al. estimate the heritability associated with common NIMs (NIMs with MAF > 5%) using stratified LD Score Regression (S-LDSR) with LD scores computed from 1KG (see **Note S2**). Because more than 70% of NIMs have MAF < 5%, this approach may not extrapolate to understand the heritability from all NIMs. An additional potential concern with analyses of NIMs is the possibility of confounding due to population structure among these introgressed variants. Typical approaches to account for population stratification based on the inclusion of principal components (PCs) may not be adequate as these PCs are computed from common SNPs on the UK Biobank genotyping array and may not account for stratification at the NIMs that tend to be rare on average (Mathieson 2012). Since our analyses work directly on individual genotype data, we are able to control for stratification specific to NIMs by including PCs estimated from NIMs in addition to PCs estimated from common SNPs. Our analyses are broadly consistent when including NIM PCs than without (see **Note S3**).

Beyond characterizing aggregate effects of NIMs, we also attempted to identify individual NIMs that modulate phenotypic variation. A challenge in identifying such variants comes from the fact that NIMs tend to have lower MAF and higher LD compared to MH SNPs. Lower MAF tends to limit the power to detect a genetic effect while higher LD makes it harder to identify the causal variant. These challenges led us to design a fine mapping strategy for prioritizing causal NIMs that enables the identification of sets of NIMs that can credibly exert influence on specific phenotypes. Using this approach, we identified credible NIMs in a number of functionally important genes, including a premature stop codon in the FCGR2A gene, and a start codon loss in COQ10A. In addition, mutations in STAT2 are found to be highly pleiotropic. As many of the genes are relevant to immune, metabolic, and developmental disorders, with functions relevant to the transition to new environments, the credible NIMs reported in our study offer a starting point for detailed investigation of the biological effects of introgressed variants. Greenbaum et al. hypothesized that introgression-based transmission of alleles related to the immune system could have helped human out-of-Africa expansion in the presence of new pathogens (Greenbaum 2019). While our results do not directly support this hypothesis, they pinpoint introgressed alleles in immune-related genes that could have and continue to modulate human phenotypes. Although we identified a number of likely causal NIMs in fine mapping, our strategy likely only picks up a small fraction of the functional NIMs suggesting that additional NIMs that are causal for specific traits remain to be discovered.

Our study has several limitations due to the current availability of data and statistical methods. First, all of our analyses focus on the white British individuals in the UKBB due to the large sample size that permits the interrogation of low-frequency NIMs and our choice of NIMs based on introgressed mutations segregating in European populations. Whole-genome sequencing data in diverse populations can potentially elucidate the impact of Neanderthal introgression in other out-of-African populations that harbor substantial Neanderthal ancestry. Alternatively, designing arrays that have SNPs informative of archaic ancestry followed by genotype imputation could be a fruitful strategy to leverage large Biobanks to systematically explore the contribution of archaic introgression. Second, while our approach to localize credible NIMs yields a list of NIMs that are highly likely to modulate variation in a trait, our method only identifies a subset of causal variants. The design of fine mapping methods to study introgressed mutations while taking into account the ancestry (as well as better incorporating other measures such as posterior inclusion probabilities) is an important direction for future work. More broadly, the unique evolutionary history of introgressed variants motivate the development of methods tailored to their population genetic properties. While our results suggest potential evolutionary models that explain our observations of depleted heritability at introgressed alleles, evolutionary models that can comprehensively explain our observations are lacking. A major challenge is the large space of potential models that need to be explored. Nevertheless, proposing and validating such models will be an important direction for future work.

## Acknowledgments

We thank Bogdan Pasaniuc and Kirk Lohmueller for feedback on the manuscript. DR was supported by an Allen Discovery Center grant on Brain Evolution from the Paul Allen Foundation, John Templeton Foundation grant 61220, US National Science Foundation HOMINID grant BCS-1032255, US National Institutes of Health grants GM100233 and HG006399, and is an Investigator of the Howard Hughes Medical Institute. SS is supported in part by NIH grants R35GM125055, NSF grants III-1705121, CAREER-1943497, an Alfred P. Sloan Research Fellowship, and a gift from the Okawa Foundation. PL was supported by a Burroughs Wellcome Fund Career Award at the Scientific Interfaces and the Next Generation Fund at the Broad Institute of MIT and Harvard.

This research was conducted using the UK Biobank Resource under applications 12408 and 33127.

## Methods

### Identification and design of SNPs that tag Neanderthal ancestry on the UK Biobank Axiom array

We chose a subset of SNPs to add to the UKBiobank Axiom array that would tag introgressed Neanderthal alleles segregating in present-day European populations.

We began with a list of 95,462 SNPs that are likely to be Neanderthal-derived from Sankararaman et al. 2014. These SNPs were identified to tag confidently inferred Neanderthal haplotypes in the European individuals identified in the 1000 Genomes Phase 1 data (**Note S1**).

We winnowed down this list to 43,026 SNPs after removing ones already tagged at *r*^2^ > 0. 8 by SNPs on the UKBiLEVE array. We then designed a greedy algorithm to capture the remaining untagged SNPs that could still be accommodated on the array (we determined the number of oligonucleotide features that would be needed to genotype each SNP as well as the total number of features available on the array through discussions with UKBiobank Axiom array design team).

Specifically, we computed LD between all pairs of Neanderthal-derived SNPs and then iteratively picked SNPs with the highest score to add to the array where the score was computed as:

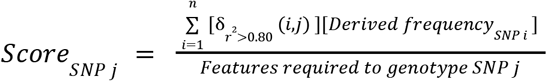

Here, 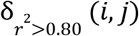 is an indicator variable that is 1 if the squared correlation coefficient between SNPs *i* and *j* is >0.80 and zero otherwise. Thus, SNP *j* is scored higher if it tags other untagged SNPs on the array. The other two terms upweight SNPs that tag other Neanderthal-derived SNPs with high derived allele frequency in Europeans and downweight SNPs by the number of oligonucleotide features required to genotype the SNP.

We iteratively chose SNPs until we obtained 6,027 SNPs (requiring 16,674 features) that fully tagged the remaining set of Neanderthal-derived SNPs. These 6,027 SNPs were then added to the UKBiobank Axiom array.

### UK Biobank (UKBB) genotype QC

We restricted all our analyses to a set of high-quality imputed SNPs (with a hard call threshold of 0.2 and an info score greater than or equal to 0.8), which, among the 291,273 imputed genotypes of UKBB unrelated white British individuals, 1) have MAF higher than 0.001,2) are under Hardy-Weinberg equilibrium (*p* > 10^-7^), and 3) are confidently imputed in more than 99% of the genomes. Additionally, we excluded SNPs in the MHC region, resulting in a total of 7,774,235 SNP which we refer to as QC-ed SNPs.

### Identification of Neanderthal Informative Mutations

We intersected the 95,462 Neanderthal-derived SNPs identified in the 1000 Genomes European individuals with UKBB QC-ed SNPs, resulting in 70,374 mutations that we term confident Neanderthal Informative Mutations (NIM). SNPs in high Linkage Disequilibrium (LD) with this set are likely introduced through Neanderthal introgression. We expanded this set by including all QC-ed SNPs, which 1) have an *r*^2^ of 0.99 or higher with any confident NIM, and 2) are located in the proximal neighborhood of any confident NIM (within 200kb). We term this set of SNPs as expanded NIMs. On average, 80.58% of expanded NIMs match the corresponding Altai Neanderthal allele, in contrast to 2.18% of the remaining SNPs, suggesting that these SNPs are also highly informative about Neanderthal ancestry. This treatment expands the number of NIMs in the UKBB QC-ed SNPs from 70,374 (confident NIMs) to 235,592 (expanded NIMs). We primarily use this more inclusive set of SNPs in our analyses, and refer to them as NIMs in the main results. SNPs that were not part of the expanded NIMs are termed modern human (MH) SNPs.

### Annotating QC-ed SNPs by MAF and LD

In addition to ancestry (Neanderthal vs MH), we annotate each QC-ed SNP by its minor allele frequency (MAF) and LD. We define five MAF-based annotations by dividing all QC-ed SNPs into five equal-sized bins by their MAFs. We similarly define five LD-based annotations by dividing all QC-ed SNPs into five equal-sized bins based on their LD-score computed from 291,273 imputed unrelated white British genotypes. In-sample LD-score is computed on QC-ed genotypes using GCTA (https://cnsgenomics.com/software/gcta/#Overview) with flags “--ld-score --ld-wind 10000”.

After each QC-ed SNP is annotated with three properties -- ancestry (NIMs vs MH), MAF, and LD, we use them to construct three additional sets of annotations: ancestry + MAF, ancestry + LD, and ancestry + MAF + LD annotations, by intersecting MAF annotation with ancestry annotation, LD annotation with ancestry annotation, and all three annotations, respectively. For example, for ancestry + MAF annotation, we intersect the previously defined MAF annotation with the ancestry annotation and divide SNPs into ten non-overlapping bins -- from low to high MAF with Neanderthal ancestry (five bins) and from low to high MAF with modern human ancestry (five bins). Similarly, when SNPs are annotated with LD + ancestry, we have five LD bins with Neanderthal ancestry corresponding to five LD groups with modern human ancestry.

Because NIMs tend to have low MAF and high LD-score (**Fig. 2**), the sizes of the annotation bins are highly uneven. To enable reliable downstream heritability analyses, we remove the annotation bins in their entirety if they include fewer than 30 SNPs. Such exceptions only occur when SNPs are annotated based on all three annotations, i.e., ancestry + MAF + LD.

### Whole-genome simulations

We simulated phenotypes based on QC-ed UKBB genotypes with the same sample size (291,273) and number of SNPs (7,774,235). In each simulation, either 10,000 variants (mimicking moderate polygenicity) or 100,000 (mimicking high polygenicity) are sampled from the QC-ed SNPs to have causal phenotypic effects while the rest of the variants have zero effect. Causal effects and phenotypes are simulated with GCTA assuming either a high SNP heritability of 0.5 or a moderate SNP heritability of 0.2.

With the simulated causal NIM variants, true NIM heritability 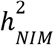 can be computed as

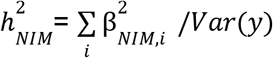

where phenotypes *y* are simulated based on a set of standardized genotype data with a simple additive genetic model

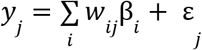

and

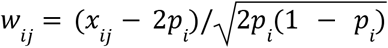

with *x_ij_* being the number of reference alleles for the *i*^th^ causal variant of the *j*^th^ individual and p. being the frequency of the *i*^th^ causal variant, *β_i_* is the allelic effect of the *i*^th^ causal variant and *ε_j_*. is the residual effect generated from a normal distribution with mean 0 and variance 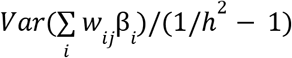.

Following previous work (Evans 2018), we chose causal variants according to five different MAF and LD-dependent genetic architecture: 1) BASELINE: baseline architecture, where SNPs are randomly selected to be causal variants, 2) COMMON: common SNPs are enriched for phenotypic effects so that SNPs with MAF > 0.05 contribute 90% of causal variants while rare SNPs contribute 10%, 3) RARE: rare variants are enriched for phenotypic effects such that SNPs with MAF <= 0.05 contribute to 90% of causal variants while the rest contribute 10%, 4) LOW: low LD SNPs are enriched for phenotypic effects, realized as SNPs whose LD-score <= 10 contribute 90% of causal variants, and the rest contribute 10%, and 5) HIGH: high LD SNPs are enriched for phenotypic effects, such that SNPs with LD-score > 10 contribute 90% causal variants while the rest contribute 10%. We simulated three replicates, for each genetic architecture with two different values of SNP heritability (0.2 and 0.5) and two different levels of polygenicity (10,000 and 100,000 causal variants). Thus, we simulated a total of 60 genetic architectures.

### Estimating NIM heritability with RHE-mc

We are interested in estimating the proportion of phenotypic variance attributed to NIMs (true NIM heritability 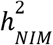) and evaluating if the heritability at a NIM (per-NIM heritability) is larger or smaller than that of a background MH SNP. To this end, we used a variance components model that partitions phenotypic variance across genomic annotations that include ancestry (NIM vs MH) as one of the input annotations.

We use RHE-mc, a method that can partition genetic variance across large sample sizes, to estimate NIM heritability (Pazokitoroudi 2020). For each phenotype, we run RHE-mc, in turn, with four types of input annotations: ancestry alone, ancestry + MAF, ancestry + LD, and ancestry + MAF + LD as described above. The ancestry+ MAF, ancestry + LD, and ancestry + MAF + LD annotations are intended to account for the differences in the MAF and LD properties of NIMs compared to MH SNPs.

To estimate NIM heritability, 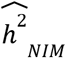, we combine the heritability of each bin corresponding to Neanderthal ancestry:

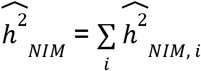

and the heritability estimates for any bins with modern human ancestry are used to compute the total heritability from MH. Thus, when we estimate NIM heritability from RHE-mc run with ancestry + MAF annotations, we add the heritability estimates from five bins of low to high MAF NIMs.

To compare the average heritability at a NIM to the heritability of a background MH SNP that is chosen to match the NIM in terms of MAF and LD profiles, we compute the following statistic:

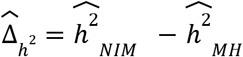

where 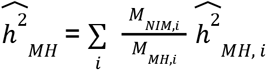 is the heritability of the background set matched for the MAF and LD profile of the set of NIMs. Here *M_MH,_i__*. denotes the number of MH SNPs in bin *i* (defined according to MAF and/or LD of the MH SNPs) while *M_NIM,_i__* denotes the number of NIMs in the corresponding bin. A more detailed justification of this statistic is provided in **Note S4**.

The standard errors (s.e.) of these statistics are computed using 100 jackknife blocks using an extension of RHE-mc that takes into account the covariance among different annotations. This new version of the RHE-mc is now available at https://github.com/alipazokit/RHEmc-coeff.

### NIM heritability and META-analysis using UKBB phenotypes

We applied RHE-mc to a total of 96 UKBBphenotypes. These phenotypes fall into 14 broader phenotypic categories (**Data S1**): anthropometry, autoimmune disorders, blood biochemistry, blood pressure, bone densitometry, environmental factors, eye, general medical information, glucose metabolism, kidney, lipid metabolism, liver, lung, and skin and hair. For each phenotype, we use RHE-mc to estimate the NIM heritability 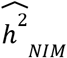 and the difference between per-NIM heritability and the per-SNP heritability of MH SNPs 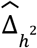 while controlling for age, sex, the first 20 genetic Principal Components (PCs) estimated from common SNPs, and the first five PCs estimated from NIMs (NIM PCs). The five NIM PCs are computed using all NIMs in unrelated white British samples with ProPCA (Agrawal 2020).

To improve power to detect patterns that are shared across groups of phenotypes, we combined analyses across groups of phenotypes and across all phenotypes analyzed. We performed random effect meta-analysis on each phenotypic category containing at least four phenotypes. We assume that the phenotypes within each category *i* have their 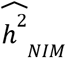 drawn from the same distribution so that we can estimate the mean 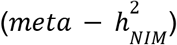 and variance of distribution *i*, based on the sampled 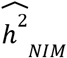 and the s.e. 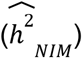. From there, we computed the meta analysis Z-score to test if the *meta* – 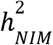 is equal to zero. Similarly, we assume the phenotypes within each category *i* have their 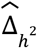 drawn from the same distribution, and compute the Z-score to test if the *meta* – Δ_*h*_2__ is equal to zero. In addition to the meta-analysis within the phenotypic category, we also performed meta-analysis across all phenotypes where we used a subset of 32 phenotypes that were chosen to have low correlation (Pearson’s *r*^2^ ≤ 0.25).

### Identifying individual NIMs associated with phenotype

To identify individual NIMs associated with a phenotype, we fit a linear regression model using plink 2.0 --glm and included covariates controlling for age, sex and the first 20 genotypic PCs, and first five NIM PCs. We used a stringent p-value threshold of 10^-10^ to correct for the number of NIMs and phenotypes tested. For each phenotype, we clumped all significant NIMs that lie within 250 kb and with an LD threshold (*r*^2^) of 0.5 using a significance threshold for the index SNP of 10^-10^.

### Identifying NIMs that modulate phenotype

To assess our ability to identify introgressed variants that truly modulate a phenotype, we first tested each NIM for association with the simulated phenotype. A challenge with such an approach is the possibility that a NIM can be found to be associated with a phenotype due to being in LD with a non-introgressed variant. To exclude settings where the association signal at a NIM might be driven by LD with a non-introgressed variant, we applied a Bayesian statistical fine-mapping method (SuSiE, https://stephenslab.github.io/susie-paper/index.html) that analyzes both NIM and MH SNPs in the region surrounding an associated NIM to output a set of SNPs that can explain the association signal at the region. Furthermore, we processed these credible sets to obtain a set of **credible NIMs**.

We performed simulations to test the accuracy of such an approach in identifying truly causal NIMs. In particular, we first ran an association test with plink (https://www.cog-genomics.org/plink/) to identify significant NIMs (p-value < 10^-10^). We then LD-pruned significant NIMs to get a subset of NIMs which are approximately uncorrelated with each other (using the plink flag “--indep-pairwise 100kb 1 0.99”). For each LD-pruned significant NIM, we considered all the QC-ed SNPs in its 200kb neighborhood as input to fine mapping. We ran SuSiE with *ρ* = 0.95 and *L* = 10, such that it returns credible sets that have at least 0.95 probability to contain one causal variant and outputs at most ten credible sets for each tested region. If there are more than one credible set for a tested region, we merge them into one set. We then removed the credible sets which have 50% or more MH SNPs in their credible set. The remaining credible sets all have majority NIMs (i.e. positive results), and they are further merged together with other such regions it overlaps with, resulting in distinct regions with evidence of NIM causal effects. We termed the set of all resulting NIMs as the **credible NIM set** and all NIMs that lie in the credible set as **credible NIMs**. The region containing the credible NIM set is termed **credible NIM region**. If there is at least one true causal NIM within the set of credible NIMs, this credible NIM region is counted as a True Positive (TP). If there is no causal NIM in the credible NIMs, this credible NIM region is counted as a False Positive (FP).

We adopted the same approach when analyzing UKBB phenotypes while incorporating covariates. Because the SuSiE package does not directly incorporate covariates, we used regression residuals from linear regression between each UKBB phenotype and UKBB covariates (age, sex, 20 regular PCs, 5 NIM PCs), as the input phenotype to SuSiE.

### Annotating NIMs

We annotated all unique credible NIMs using SnpEff (Cingolani 2012) which uses Sequence Ontology (http://www.sequenceontology.org/) to assign standardized terminology for assessing sequence change and impact. We primarily focused on examining the high (e.g., start codon loss, stop codon gain) and moderate impact SNPs (nonsynonymous variants) which are coding variants that alter protein sequences.

## Supplementary Figures

### Include Supplementary Figure S1-S7

**Supplementary Figure S1.**
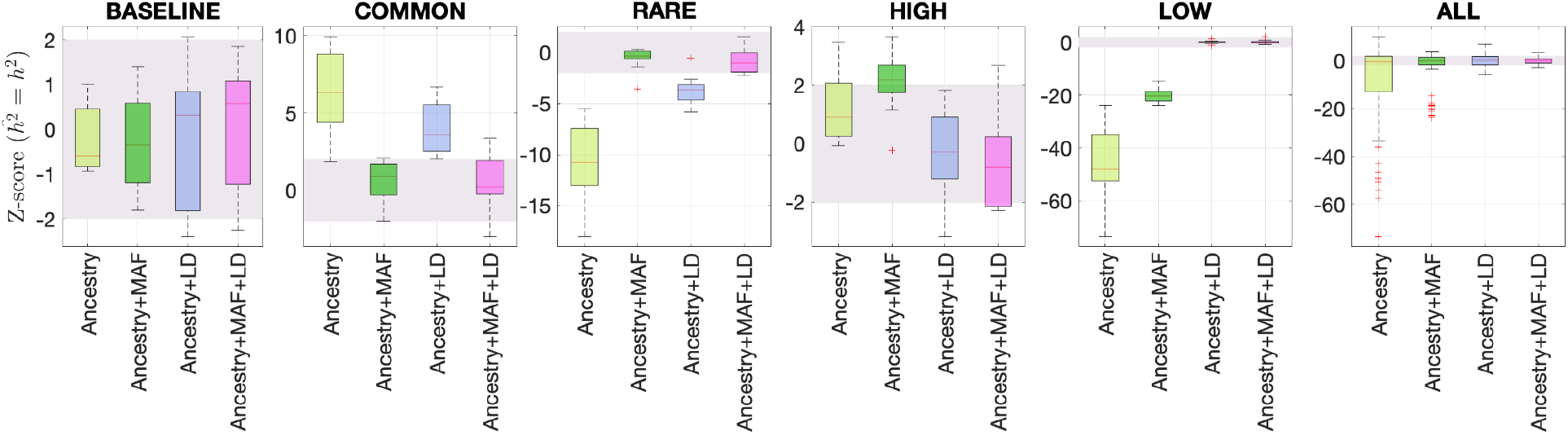
Benchmarking different methods for estimating the total SNP heritability. We grouped the simulations by the five different MAF-LD coupling: BASELINE, COMMON, RARE, HIGH, LOW, as labeled on top of each column. In each group, there are 12 simulations with different levels of polygenicity and heritability (see **Methods**). Additionally, we combined simulations from all five architectures together as ALL for the sixth column. On the y-axis, Z-score 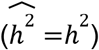 tests whether the estimated and simulated total heritability are equal. In each panel, the results from RHE-mc with four different annotations, ancestry only, ancestry + MAF, ancestry + LD, ancestry + MAF + LD are shown on the x-axis. A calibrated method is expected to have all Z-scores distributed around zero and within ±2 (shaded region). Among all tested methods, only RHE-mc with annotation ancestry + MAF + LD satisfies this criterion.

**Supplementary Figure S2.**
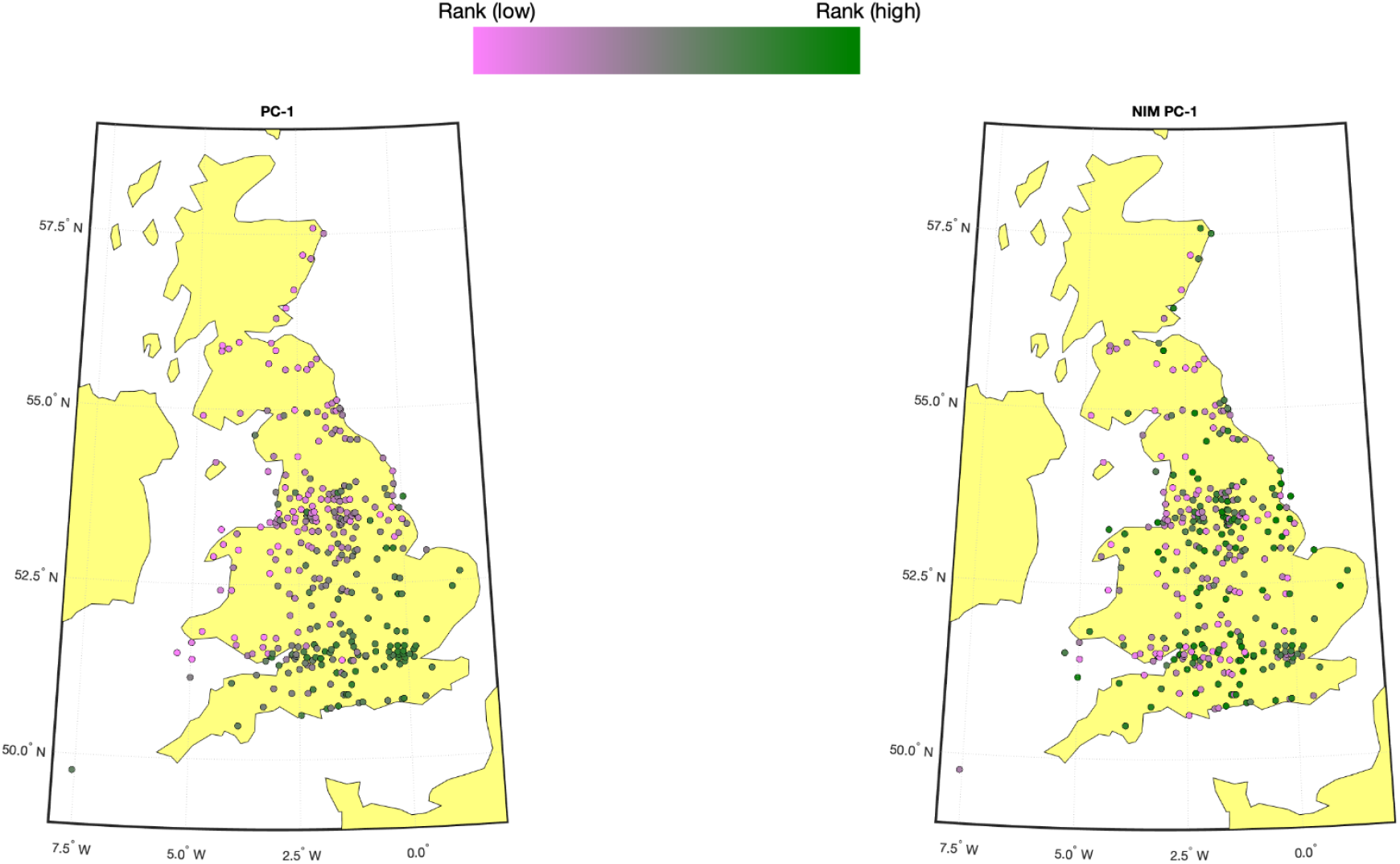
Population structure within white British samples. PC-1 from the whole genome genotypes (released by UKBB) is shown on the left, and NIM PC-1 is shown on the right. We used a 20-by-20 grid along the latitude and longitude, dividing the map into 400 colonies. We then computed the average PC projection as well as the median longitude and latitude among the individuals belonging to each colony, if there are at least 10 individuals in a colony. Each color-filled circle with a 5 kilometer radius represents one colony on the map. To maximize the visible differences, we sorted the colonies by their PC values and used the rank to determine the color of the colony. Compared to NIM PC-1, PC-1 shows a much stronger correlation with geographical location.

**Supplementary Figure S3.**
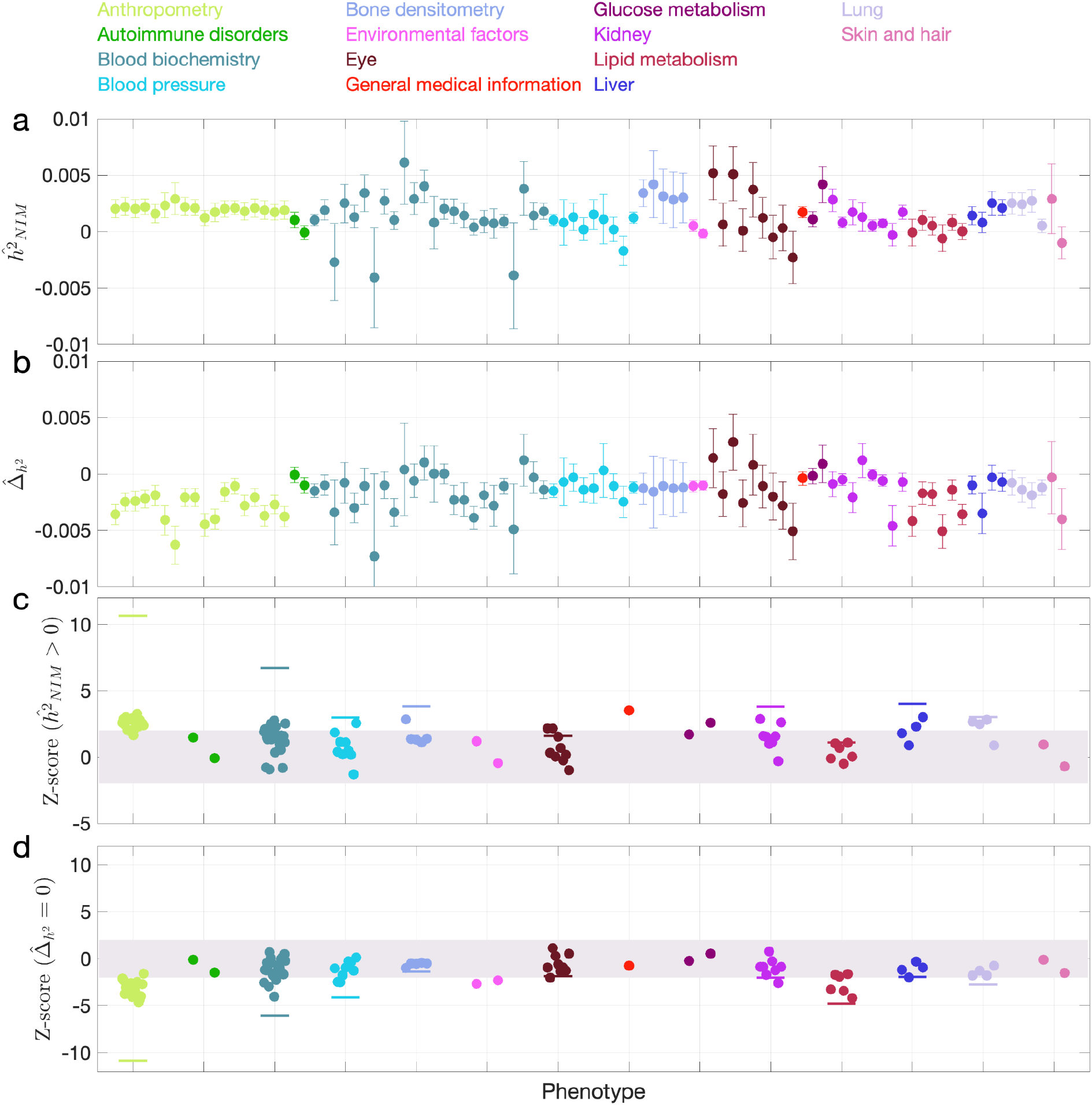
NIM heritability in the 96 UKBB phenotypes without controlling for NIM PCs. This figure is plotted in the same way as **Fig. 3**. Heritability estimates are largely similar, but fewer phenotypes are significant. Three phenotypes have significant positive NIM heritability (Z-score 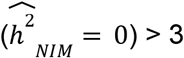): overall health rating, waist-hip-ratio, and gamma glutamyltransferase. Fourteen phenotypes (standing height, sitting height, weight, body fat percentage, whole body fat-free mass, whole body water mass, trunk fat-free mass, trunk predicted mass, basal metabolic rate, RBC count, apolipoprotein A, HDL cholesterol, triglycerides) are significantly depleted for NIM heritability (Z-score < −3).

**Supplementary Figure S4.**
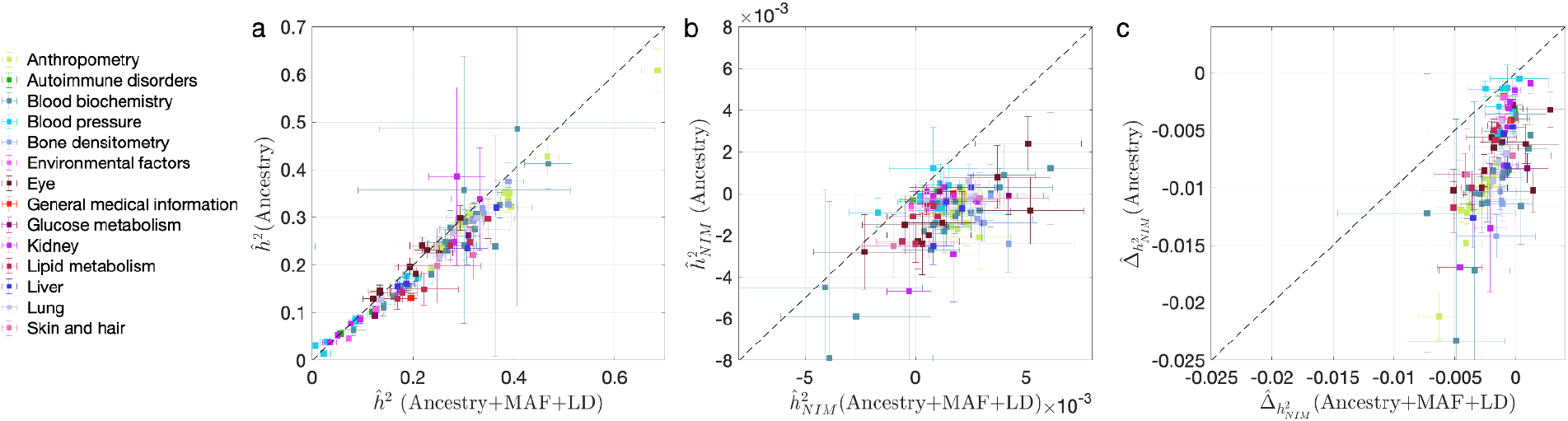
Comparing heritability estimates from RHE-mc without controlling for NIM PCs with Ancestry+MAF+LD annotation and RHE-mc with Ancestry annotation in UKBB phenotypes. This figure is plotted in the same way as **Fig. 4**. The trend that not controlling for MAF and LD lead to underestimation of **(a)** total heritability 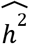, **(b)** NIM heritability 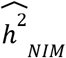, and stronger NIM heritability depletion **(c)** 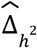 is also apparent when NIM PCs are not controlled for.

**Supplementary Figure S5.**
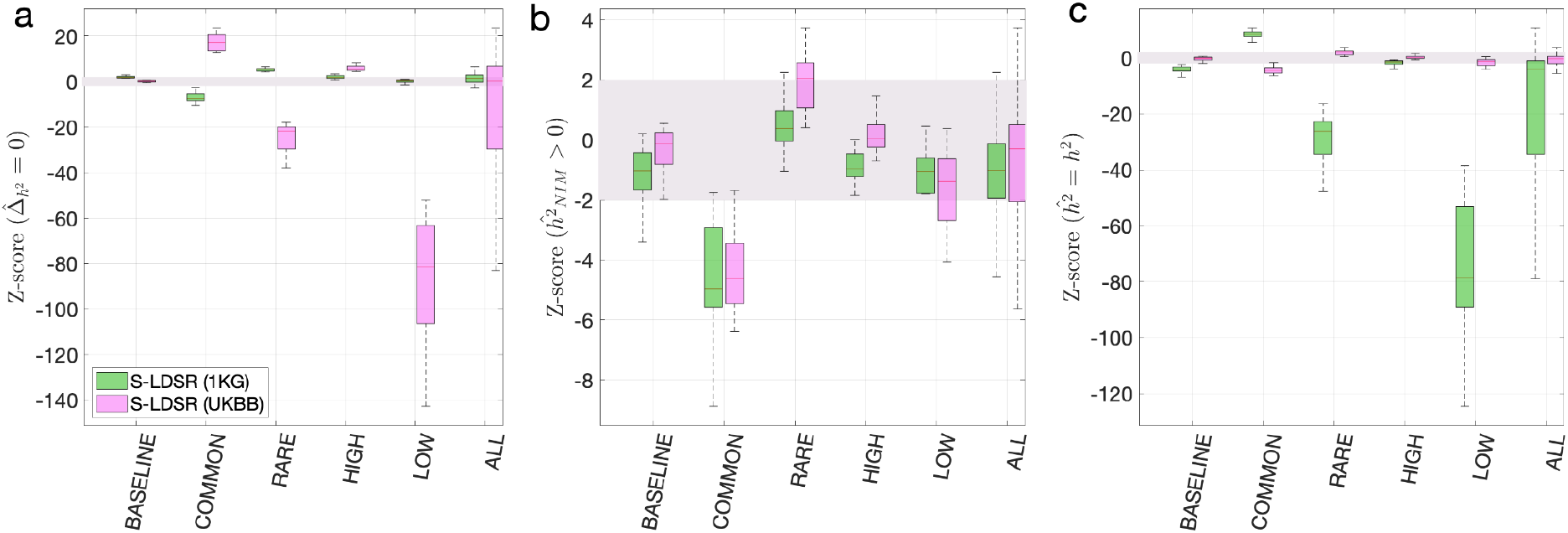
Benchmark stratified LDSC regression (S-LDSR) with in-sample and out-of-sample LD scores. We group the simulations by the MAF-LD coupling: BASELINE, COMMON, RARE, HIGH, LOW, and ALL, as labeled on the x-axis. We plot the distributions of three Z-scores (y-axis), one on each panel: **(a)** Z-score 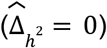 tests whether the estimated NIM heritability is different from the matched MH heritability, **(b)** Z-score 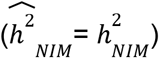 tests whether the estimated and expected NIM heritability are equal, and **(c)** Z-score 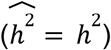 tests whether the estimated and simulated total heritability are equal. In each panel, S-LDSR with the out-of-sample LD score from 1000 Genomes (1KG) is shown in green and S-LDSR with in-sample LD score from UKBB in pink. In S-LDSR, only ancestry annotation is used. The Z-scores within ±2 are color shaded. S-LDSR (1KG) is not calibrated even for BASELINE architecture.

**Supplementary Figure S6.**
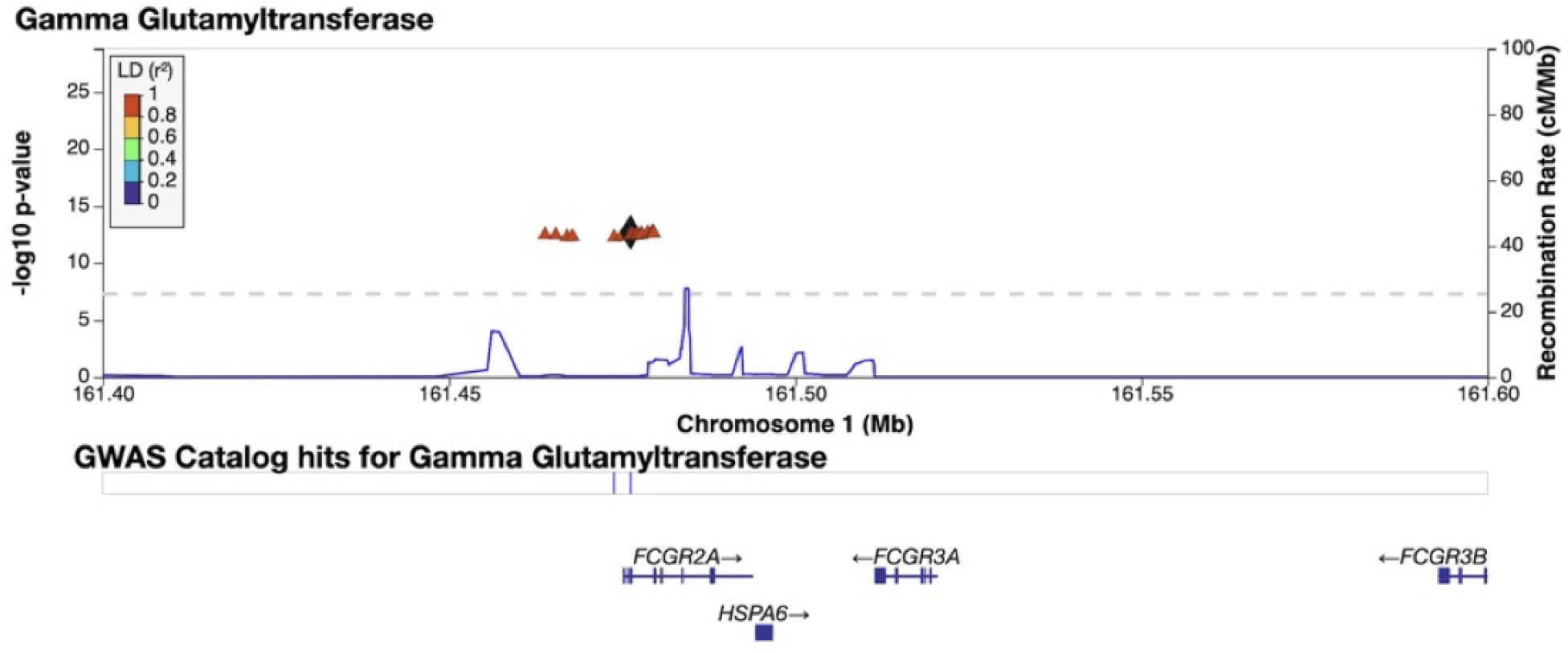
Credible NIM in the FCGR2A gene associated with gamma-glutamyl transferase levels. Plot of 200kb region surrounding rs9427397 (marked in black diamond; hg19 coordinates), a credible NIM in FCGR2A that introduces a premature stop codon and is associated with increased levels of gamma glutamyltransferase (while also associated with increased levels of aspartate aminotransferase and decreased total protein). The plot displays other NIMs in the region along with their LD (r^2^) to rs9427397 computed in 1000 Genomes Europeans.

**Supplementary Figure S7.**
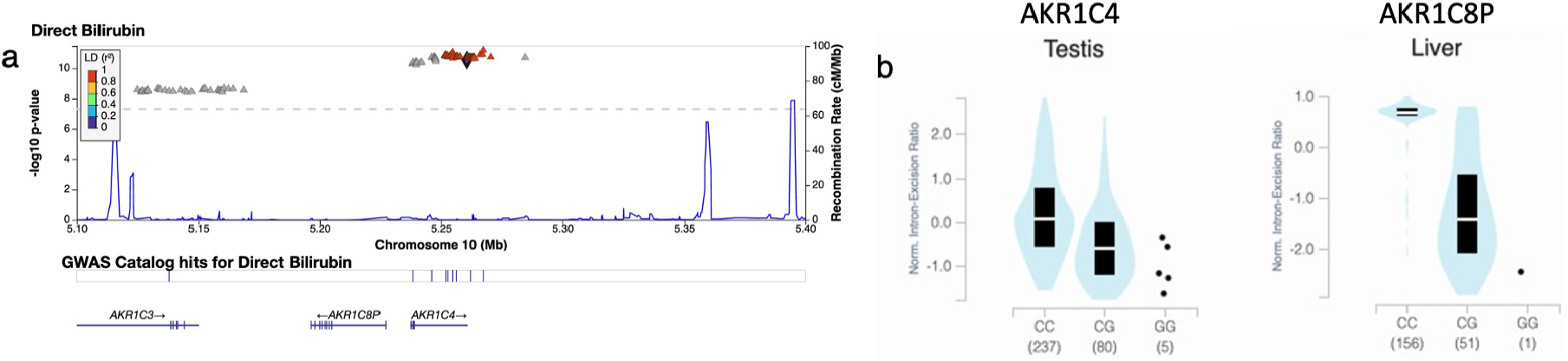
Credible NIM in the AKR1C4 gene is associated with bilirubin levels. **(a)** Plot of 300kb region surrounding rs17134592 (marked in black diamond; hg19 coordinates), a non-synonymous NIM in AKR1C4, that is associated with increased serum bilirubin levels. The plot displays other NIMs in the region along with their LD (*r*^2^) to rs17134592. **(b)** rs17134592 is a splicing QTL in liver (AKR1C8P) and testis (AKR1C4) identified in GTEx v8.

## Supplementary data

Supplementary data include 10 items, Data S1-S10 are available at https://github.com/AprilWei001/NIM

1. **Data S1**. UKBB phenotype annotation.
2. **Data S2**: RHE-mc results in simulated data.
3. **Data S3**: RHE-mc results with Ancestry+MAF+LD annotations and NIM PCs included in covariates applied to 96 UKBB phenotypes
4. **Data S4**: RHE-mc results with Ancestry only annotation and NIM PCs included in covariates applied to 96 UKBB phenotypes
5. **Data S5**: RHE-mc results with Ancestry+MAF+LD annotation without NIM PC in covariates applied to 96 UKBB phenotypes
6. **Data S6**: Fine mapping FDP in simulated data.
7. **Data S7:** 112 credible NIM sets and credible NIMs
8. **Data S8**: SnpEff annotation of all unique credible NIMs
9. **Data S9.** Stratified LD score regression results in simulated data using LD score from 1KG
10. **Data S10.** Stratified LD score regression results in simulated data using LD score from UKBB

## Supplementary Notes

### Supplementary Note S1: Identification of SNPs that tag Neanderthal ancestry on the UK Biobank Axiom array

Starting with the confidently inferred Neanderthal haplotypes identified in Sankararaman et al. 2014, we identified whether a SNP segregating in a target modern human population owes its origin to the Neanderthal gene flow event as follows:

1. We identified sets of haplotypes that are confidently labeled as Neanderthal, *N* by the Conditional Random Field (CRF) method proposed in Sankararaman et al. 2014 scanning for runs of consecutive SNPs with marginal probability of Neanderthal ancestry ≥0.90. We also identified sets of haplotypes that are confidently labeled as non-Neanderthal, *MH* by scanning for SNPs with marginal probability ≤0.1). We also required the Neanderthal haplotype to be at least 0.02 cM long.
2. For each SNP called in the 1000 Genomes dataset, we required that none of the derived alleles at this SNP falls on one of the modern human haplotypes in the set *MH* and all of the haplotypes in *N* carry the derived allele. This procedure allows for some false negatives in the predictions of the CRF.
3. We ran this procedure on the combined calls from the European ancestry populations (CEU, GBR, FIN, IBS and TSI) in the 1000 Genomes Project.

This procedure yielded a total of 95,462 SNPs that are likely to be Neanderthal-derived. We winnowed down this list to 43,026 SNPs after removing ones already tagged at *r*^2^ > 0.8 by SNPs on the UKBiLEVE array. We then designed a greedy algorithm to capture the remaining untagged SNPs that could still be accommodated on the array (we determined the number of oligonucleotide features that would be needed to genotype each SNP as well as the total number of features available on the array through discussions with UKBiobank Axiom array design team).

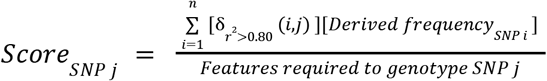

Here 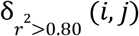 is an indicator variable that is 1 if the squared correlation coefficient between SNPs *i* and *j* is >0.80 and zero otherwise. Thus, SNP j is scored higher if it tags other untagged SNPs on the array. The other two terms upweight SNPs that tag other SNPs with high derived allele frequency in Europeans and downweight SNPs by the number of oligonucleotide features required to genotype it.

### Supplementary Note S2: Estimating NIM heritability with partitioned LD-score regression

We considered two candidate methods for estimating the NIM heritability in large datasets and testing the related hypotheses to NIM heritability, S-LDSR (Finucane 2015) and RHE-mc (see Main text) (Pazokitoroudi 2020). S-LDSR can speedily estimate partitioned heritability given GWAS statistics and LD scores without any individual-level data. S-LDSR can be used with either in-sample LD scores (i.e., computed from the same data as for GWAS) or out-of sample LD scores (i.e., computed from an external and often much smaller data set). Out-of-sample LD scores from 1000 Genomes (1KG) is often used in S-LDSR (McArthur 2021, Koller 2021) because 1) it is computationally much cheaper to compute than using the GWAS cohorts, and 2) individual-level data from GWAS cohorts are not always accessible; despite that, S-LDSR with in-sample LD scores is more accurate in theory.

Previous studies by Koller et al and McArthur et al used S-LDSR to estimate the heritability from archaic ancestries. They computed the stratified LD scores using the 1000 Genomes (1KG) EUR and EAS samples and performed LD score regression against the GWAS statistics from a different cohort. If the ancestry from 1KG samples does not match well with the GWAS cohort, it could lead to biased heritability estimates. Additionally, the LD score distribution and MAF distributions of NIMs are very different from the distributions of MH SNPs (**Fig. 1**), which might also affect the heritability estimates if not taken into account. Finally, LD score regression is restricted to a subset of SNPs (typically with MAF > 5%) which substantially reduces the number of NIMs analyzed. Here, we benchmarked S-LDSR on the simulated data with both out-of-sample LD scores from 1KG and the in-sample LD scores from all UKBB QC-ed data, stratified by ancestry (NIM vs MH).

First, we used the aforementioned simulations to evaluate the partitioned LD score regression in estimating NIM heritability. We downloaded the 1KG EUR data (from https://storage.googleapis.com/broad-alkesgroup-public/LDSCORE/1000G_Phase3_plinkfiles.tgz) that is typically used for LD score regression. There are 9,997,231 SNPs in the data, and 5,789,471 of them are shared with the UKBB QC-ed SNPs. Out of the 235,592 expanded NIMs defined in UKBB QC-ed data, 210,962 are present in 1KG EUR, and we refer to these SNPs as the 1KG NIMs. We defined the 9,786,269 SNPs in 1KG EUR that are not expanded NIMs as 1KG MH SNPs. We then computed the stratified LD score using GCTA software with flags --ld-score --ld-wind 10000 for all the 1KG SNPs, all the 1KG NIMs, and all the 1KG MH SNPs. From here, we computed the stratified LD score with *Idsc_SNP_* = *Idsc_NIM_* + *Idsc_MH’_*, such that each SNP has two stratified LD scores, one due to its LD with NIMs and another due to its LD with MH SNPs. We then intersected the 1KG SNPs with UKBB QC-ed SNPs, and used the shared 5,789,471 SNPs to perform stratified LD score regression. The S-LDSR is then performed with all SNPs that overlap between 1KG and UKBB. For each simulation, we ran S-LDSR to estimate 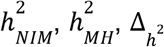, and their standard errors from 200 jackknife blocks. We found that the results from using out-of-sample LD are biased even when heritability does not depend on MAF and LD (i.e., BASELINE) (**Figure S5**).

As a comparison, we computed the in-sample stratified LD score using the UKBB QC-ed data and applied S-LDSR with these in-sample LD scores. In contrast to the previous results, the results are well calibrated for BASELINE, suggesting that the previous biases observed with BASELINE are due to the disagreement between the out-of-sample LD score and the in-sample LD score (**Figure S5**). Not surprisingly, the results for MAF and LD-dependent architectures are still biased, as these factors are not taken into account. We caution that our simulations are based on UKBB QC-ed SNPs, where non QC-ed SNPs do not have an impact on the simulated phenotypes. This setting will favor S-LDSR based on UKBB QC-ed SNPs more than in actual settings, and disfavor S-LDSR 1KG more than in actual settings. It is possible that in reality, the biases with in-sample LD score will become larger, and the biases with out-of-sample LD score will become smaller. Nonetheless, because it is often expensive to compute in-sample LD scores, the accuracy will largely depend on how well the external panel resembles the GWAS cohort.

The out-of-sample LD score could be particularly biased for low MAF SNPs, hence S-LDSR recommends not using annotations with fewer than 5% of SNPs as best practice. This practice will necessarily exclude more than 70% of NIMs and about half of the MH SNPs, and the heritability estimates from high MAF SNPs may not extrapolate to low MAF SNPs. Therefore, S-LDSR, under the best practice, is not suitable for studying Neanderthal introgressed variants.

## Supplementary Note S3: The impact of inclusion of NIM PCs on NIM heritability estimates

We computed the first five NIM PCs using all NIMs in unrelated white British samples with ProPCA (Agrawal 2020). Compared to the regular genetic PCs (estimated from common SNPs), NIM PCs are only weakly correlated with birth GPS locations (**Fig. S2**), consistent with the fact that Neanderthal introgression occurred soon after the out-of-African migration before population expansion.

When NIM PCs were not being controlled for (with remaining regular covariates still used), we found three phenotypes with significant NIM heritability (Z-score 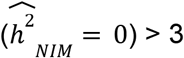): overall health rating, waist-hip-ratio (WHR), and gamma glutamyltransferase (a measure of liver function). We also combined phenotypes into broader phenotypic categories and performed random effect meta-analysis on the nine categories that contain at least four phenotypes (see **Methods**). We found that meta – 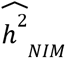 is significantly larger than zero (Z-score > 2.53 for one-tail *p* = 0.05 level) for all but two categories (eye, lipid metabolism), meaning that NIMs heritability is generally nonzero (**Fig. S3ac**). We then tested whether NIM heritability is larger or smaller compared to MH SNPs 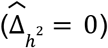. Fourteen phenotypes (standing height, sitting height, weight, body fat percentage, whole body fat-free mass, whole body water mass, trunk fat-free mass, trunk predicted mass, basal metabolic rate, RBC count, apolipoprotein A, HDL cholesterol, triglycerides) remain significantly depleted (Z-score < −3), among which eight are anthropometric phenotypes, and three are related to lipid metabolism. This is in contrast to seventeen phenotypes when NIM PCs are controlled for (body mass index, hip circumference, waist circumference, standing height, sitting height, weight, whole body fat-free mass, whole body water mass, whole body impedance, trunk fat mass, trunk fat-free mass, trunk predicted mass, basal metabolic rate, RBC count, apolipoprotein A, HDL cholesterol, triglycerides).

Four phenotypic categories show significant NIM heritability depletion (anthropometry, blood biochemistry, blood pressure, lipid metabolism), and five are not significantly different with meta analysis (**Fig. S3bd**). In contrast to the evidence for depletion in NIM heritability, we found no evidence for traits with elevated NIM heritability even when excluding NIM PCs (**Fig S4c**).

## Supplementary Note S4: Statistic to compare per-NIM heritability to per-SNP heritability at a set of background MH SNPs

In this note, we provide additional intuition behind our statistic to compare difference between the heritability at a NIM (per-NIM heritability) and the per-SNP heritability of a background set of MH SNPs:

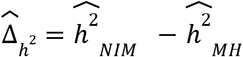

Let 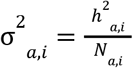 where *a* ∈ {*NIM, MH*}, *i* denotes one of the annotations (MAF,LD), 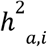 denotes the heritability attributed to annotation (*a, i*), and *M_a,i_* denotes the number of SNPs in annotation (*a, i*). Thus 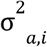 denotes the per-SNP heritability associated with annotation (*a, i*).

The per-SNP heritability associated with NIMs is given by:

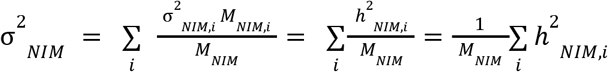

where *M_NIM_* denotes the total number of NIMs.

To choose a background set of MH SNPs that match the NIMs in terms of their MAF and LD distribution, we would pick a given bin *i* with probability 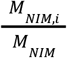. The per-SNP heritability associated with this background set of MH SNPs is then given by:

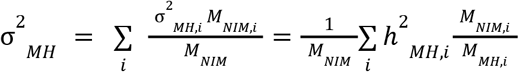

Thus, we are interested in testing the null hypothesis that the per-NIM heritability is equal to the per-SNP heritability of the background set of MH SNPs.

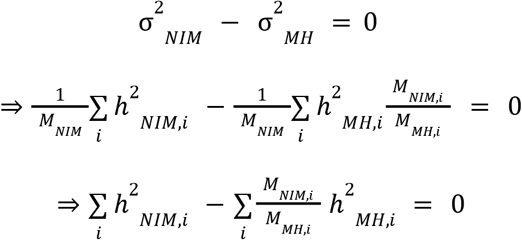

Defining our parameter of interest 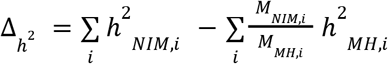, our null hypothesis is that Δ_*h*^2^_ = 0.

We estimate the relative reduction in NIM heritability as:

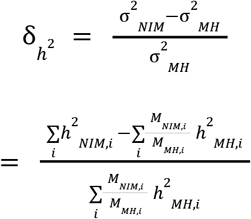

## Notes

https://github.com/AprilWei001/NIM

### Competing Interest Statement

The authors have declared no competing interest.

## References

1. M. Dannemann, J. Kelso, The Contribution of Neanderthals to Phenotypic Variation in Modern Humans. The American Journal of Human Genetics. 101, 578–589 (2017).

2. L. Skov, M. Coll Macià, G. Sveinbjörnsson, F. Mafessoni, E. A. Lucotte, M. S. Einarsdóttir, H. Jonsson, B. Halldorsson, D. F. Gudbjartsson, A. Helgason, M. H. Schierup, K. Stefansson, The nature of Neanderthal introgression revealed by 27,566 Icelandic genomes. Nature. 582, 78–83 (2020).

3. E. McArthur, D. C. Rinker, J. A. Capra, Quantifying the contribution of Neanderthal introgression to the heritability of complex traits. Nat Commun. 12, 4481 (2021).

4. C. N. Simonti, B. Vernot, L. Bastarache, E. Bottinger, D. S. Carrell, R. L. Chisholm, D. R. Crosslin, S. J. Hebbring, G. P. Jarvik, I. J. Kullo, R. Li, J. Pathak, M. D. Ritchie, D. M. Roden, S. S. Verma, G. Tromp, J. D. Prato, W. S. Bush, J. M. Akey, J. C. Denny, J. A. Capra, The phenotypic legacy of admixture between modern humans and Neandertals. Science. 351, 737–741 (2016).

5. S. Sankararaman, S. Mallick, M. Dannemann, K. Prüfer, J. Kelso, S. Pääbo, N. Patterson, D. Reich, The genomic landscape of Neanderthal ancestry in present-day humans. Nature. 507, 354–357 (2014).

6. A. Pazokitoroudi, Y. Wu, K. S. Burch, K. Hou, A. Zhou, B. Pasaniuc, S. Sankararaman, Efficient variance components analysis across millions of genomes. Nat Commun. 11, 4020 (2020).

7. I. Mathieson, G. McVean, Differential confounding of rare and common variants in spatially structured populations. Nat Genet. 44, 243–246 (2012).

8. K. Harris, R. Nielsen, The Genetic Cost of Neanderthal Introgression. Genetics. 203, 881–891 (2016).

9. G. Wang, A. Sarkar, P. Carbonetto, M. Stephens, A simple new approach to variable selection in regression, with application to genetic fine mapping. J. R. Stat. Soc. B. 82, 1273–1300 (2020).

10. P. Cingolani, A. Platts, L. L. Wang, M. Coon, T. Nguyen, L. Wang, S. J. Land, X. Lu, D. M. Ruden, A program for annotating and predicting the effects of single nucleotide polymorphisms, SnpEff: SNPs in the genome of Drosophila melanogaster strain w^1118^; iso-2; iso-3. Fly. 6, 80–92 (2012).

11. V. A. Laufer, H. K. Tiwari, R. J. Reynolds, M. I. Danila, J. Wang, J. C. Edberg, R. P. Kimberly, L. C. Kottyan, J. B. Harley, T. R. Mikuls, P. K. Gregersen, D. M. Absher, C. D. Langefeld, D. K. Arnett, S. L. Bridges, Jr, Genetic influences on susceptibility to rheumatoid arthritis in African-Americans. Human Molecular Genetics. 28, 858–874 (2019).

12. C. M. Quinzii, L. C. López, A. Naini, S. DiMauro, M. Hirano, Human CoQ _10_ deficiencies. BioFactors. 32, 113–118 (2008).

13. N. Kubota, M. Suyama, An integrated analysis of public genomic data unveils a possible functional mechanism of psoriasis risk via a long-range ERRFI1 enhancer. BMC Med Genomics. 13, 8 (2020).

14. F. L. Mendez, J. C. Watkins, M. F. Hammer, A Haplotype at STAT2 Introgressed from Neanderthals and Serves as a Candidate of Positive Selection in Papua New Guinea. The American Journal of Human Genetics. 91, 265–274 (2012).

15. T. Kume, H. Iwasa, H. Shiraishi, T. Yokoi, K. Nagashima, M. Otsuka, T. Terada, T. Takagi, A. Hara, T. Kamataki, Characterization of a novel variant (S145C/L311V) of 3alpha-hydroxysteroid/dihydrodiol dehydrogenase in human liver. Pharmacogenetics. 9, 763–771 (1999).

16. L. M. Evans, R. Tahmasbi, S. I. Vrieze, G. R. Abecasis, S. Das, S. Gazal, D. W. Bjelland, T. R. de Candia, Haplotype Reference Consortium, M. E. Goddard, B. M. Neale, J. Yang, P. M. Visscher, M. C. Keller, Comparison of methods that use whole genome data to estimate the heritability and genetic architecture of complex traits. Nat Genet. 50, 737–745 (2018).

17. Y. Okada, D. Wu, G. Trynka, T. Raj, C. Terao, K. Ikari, Y. Kochi, K. Ohmura, A. Suzuki, S. Yoshida, R. R. Graham, A. Manoharan, W. Ortmann, T. Bhangale, J. C. Denny, R. J. Carroll, A. E. Eyler, J. D. Greenberg, J. M. Kremer, D. A. Pappas, L. Jiang, J. Yin, L. Ye, D.-F. Su, J. Yang, G. Xie, E. Keystone, H.-J. Westra, T. Esko, A. Metspalu, X. Zhou, N. Gupta, D. Mirel, E. A. Stahl, D. Diogo, J. Cui, K. Liao, M. H. Guo, K. Myouzen, T. Kawaguchi, M. J. H. Coenen, P. L. C. M. van Riel, M. A. F. J. van de Laar, H.-J. Guchelaar, T. W. J. Huizinga, P. Dieudé, X. Mariette, S. L. Bridges, A. Zhernakova, R. E. M. Toes, P. P. Tak, C. Miceli-Richard, S.-Y. Bang, H.-S. Lee, J. Martin, M. A. Gonzalez-Gay, L. Rodriguez-Rodriguez, S. Rantapää-Dahlqvist, L. Arlestig, H. K. Choi, Y. Kamatani, P. Galan, M. Lathrop, RACI consortium, GARNET consortium, S. Eyre, J. Bowes, A. Barton, N. de Vries, L. W. Moreland, L. A. Criswell, E. W. Karlson, A. Taniguchi, R. Yamada, M. Kubo, J. S. Liu, S.-C. Bae, J. Worthington, L. Padyukov, L. Klareskog, P. K. Gregersen, S. Raychaudhuri, B. E. Stranger, P. L. De Jager, L. Franke, P. M. Visscher, M. A. Brown, H. Yamanaka, T. Mimori, A. Takahashi, H. Xu, T. W. Behrens, K. A. Siminovitch, S. Momohara, F. Matsuda, K. Yamamoto, R. M. Plenge, Genetics of rheumatoid arthritis contributes to biology and drug discovery. Nature. 506, 376–381 (2014).

18. A. Agrawal, A. M. Chiu, M. Le, E. Halperin, S. Sankararaman, Scalable probabilistic PCA for large-scale genetic variation data. PLoS Genet. 16, e1008773 (2020).

19. B. Vernot, J. M. Akey, Resurrecting Surviving Neandertal Lineages from Modern Human Genomes. Science. 343, 1017–1021 (2014).

20. F. Racimo, S. Sankararaman, R. Nielsen, E. Huerta-Sánchez, Evidence for archaic adaptive introgression in humans. Nat Rev Genet. 16, 359–371 (2015).

21. R. M. Gittelman, J. G. Schraiber, B. Vernot, C. Mikacenic, M. M. Wurfel, J. M. Akey, Archaic Hominin Admixture Facilitated Adaptation to Out-of-Africa Environments. Current Biology. 26, 3375–3382 (2016).

22. I. Juric, S. Aeschbacher, G. Coop, The Strength of Selection against Neanderthal Introgression. PLoS Genet. 12, e1006340 (2016).

23. M. Petr, S. Pääbo, J. Kelso, B. Vernot, Limits of long-term selection against Neandertal introgression. Proc Natl Acad Sci USA. 116, 1639–1644 (2019).

24. L. Abi-Rached, M. J. Jobin, S. Kulkarni, A. McWhinnie, K. Dalva, L. Gragert, F. Babrzadeh, B. Gharizadeh, M. Luo, F. A. Plummer, J. Kimani, M. Carrington, D. Middleton, R. Rajalingam, M. Beksac, S. G. E. Marsh, M. Maiers, L. A. Guethlein, S. Tavoularis, A.-M. Little, R. E. Green, P. J. Norman, P. Parham, The Shaping of Modern Human Immune Systems by Multiregional Admixture with Archaic Humans. Science. 334, 89–94 (2011).

25. C. Bycroft, C. Freeman, D. Petkova, G. Band, L. T. Elliott, K. Sharp, A. Motyer, D. Vukcevic, O. Delaneau, J. O’Connell, A. Cortes, S. Welsh, A. Young, M. Effingham, G. McVean, S. Leslie, N. Allen, P. Donnelly, J. Marchini, The UK Biobank resource with deep phenotyping and genomic data. Nature. 562, 203–209 (2018).

26. A. P. Boughton, R. P. Welch, M. Flickinger, P. VandeHaar, D. Taliun, G. R. Abecasis, M. Boehnke, LocusZoom.js: interactive and embeddable visualization of genetic association study results. Bioinformatics. 37, 3017–3018 (2021).

27. K. Prüfer, F. Racimo, N. Patterson, F. Jay, S. Sankararaman, S. Sawyer, A. Heinze, G. Renaud, P. H. Sudmant, C. de Filippo, H. Li, S. Mallick, M. Dannemann, Q. Fu, M. Kircher, M. Kuhlwilm, M. Lachmann, M. Meyer, M. Ongyerth, M. Siebauer, C. Theunert, A. Tandon, P. Moorjani, J. Pickrell, J. C. Mullikin, S. H. Vohr, R. E. Green, I. Hellmann, P. L. F. Johnson, H. Blanche, H. Cann, J. O. Kitzman, J. Shendure, E. E. Eichler, E. S. Lein, T. E. Bakken, L. V. Golovanova, V. B. Doronichev, M. V. Shunkov, A. P. Derevianko, B. Viola, M. Slatkin, D. Reich, J. Kelso, S. Pääbo, The complete genome sequence of a Neanderthal from the Altai Mountains. Nature. 505, 43–49 (2014).

28. R. E. Green, J. Krause, A. W. Briggs, T. Maricic, U. Stenzel, M. Kircher, N. Patterson, H. Li, W. Zhai, M. H.-Y. Fritz, N. F. Hansen, E. Y. Durand, A.-S. Malaspinas, J. D. Jensen, T. Marques-Bonet, C. Alkan, K. Prüfer, M. Meyer, H. A. Burbano, J. M. Good, R. Schultz, A. Aximu-Petri, A. Butthof, B. Höber, B. Höffner, M. Siegemund, A. Weihmann, C. Nusbaum, E. S. Lander, C. Russ, N. Novod, J. Affourtit, M. Egholm, C. Verna, P. Rudan, D. Brajkovic, Ž. Kucan, I. Gušic, V. B. Doronichev, L. V. Golovanova, C. Lalueza-Fox, M. de la Rasilla, J. Fortea, A. Rosas, R. W. Schmitz, P. L. F. Johnson, E. E. Eichler, D. Falush, E. Birney, J. C. Mullikin, M. Slatkin, R. Nielsen, J. Kelso, M. Lachmann, D. Reich, S. Pääbo, A Draft Sequence of the Neandertal Genome. Science. 328, 710–722 (2010).

29. G. Greenbaum, W. M. Getz, N. A. Rosenberg, M. W. Feldman, E. Hovers, O. Kolodny, Disease transmission and introgression can explain the long-lasting contact zone of modern humans and Neanderthals. Nat Commun. 10, 5003 (2019).

## References

1. A. Pazokitoroudi, Y. Wu, K. S. Burch, K. Hou, A. Zhou, B. Pasaniuc, S. Sankararaman, Efficient variance components analysis across millions of genomes. Nat Commun. 11, 4020 (2020).

2. H. K. Finucane, B. Bulik-Sullivan, A. Gusev, G. Trynka, Y. Reshef, P.-R. Loh, V. Anttila, H. Xu, C. Zang, K. Farh, S. Ripke, F. R. Day, ReproGen Consortium, Schizophrenia Working Group of the Psychiatric Genomics Consortium, RACI Consortium, S. Purcell, E. Stahl, S. Lindstrom, J. R. B. Perry, Y. Okada, S. Raychaudhuri, M. J. Daly, N. Patterson, B. M. Neale, A. L. Price, Partitioning heritability by functional annotation using genome-wide association summary statistics. Nat Genet. 47, 1228–1235 (2015).

4. D. Koller, F. R. Wendt, G. A. Pathak, A. De Lillo, F. De Angelis, B. Cabrera-Mendoza, S. Tucci, R. Polimanti, “The impact of evolutionary processes in shaping the genetics of complex traits in East Asia and Europe: a specific contribution from Denisovan and Neanderthal introgression” (preprint, Genetics, 2021),, doi:10.1101/2021.08.12.456138.

## References

1. A. Agrawal, A. M. Chiu, M. Le, E. Halperin, S. Sankararaman, Scalable probabilistic PCA for large-scale genetic variation data. PLoS Genet. 16, e1008773 (2020).

